# A model of apical-out human first trimester trophoblast organoids to study regeneration after syncytial damage

**DOI:** 10.1101/2025.10.21.683405

**Authors:** Ida Calvi, Giada Guntri, Alexandra Graff Meyer, Lhéanna Klaeylé, Qian Li, Jan Seebacher, Hans-Rudolf Hotz, Naomi McGovern, Margherita Y. Turco

## Abstract

The villous syncytiotrophoblast in humans is the interface between maternal blood and the placenta and its functions are essential for a successful pregnancy. The syncytium can be damaged *in vivo* focally by high-velocity and turbulent maternal blood flow, or more globally by oxidative stress. It then quickly regenerates, a process critical for the maintenance of placental functions including nutrient transfer and protection from infections. To investigate this process, we developed apical-out trophoblast organoids from first-trimester placentas and induced mechanical damage to the syncytium. We observed spontaneous regeneration of syncytiotrophoblast both morphologically and functionally. TNFRSF12A, the TWEAK receptor, was identified as a potential enhancer of syncytiotrophoblast regeneration. Supplementation with recombinant TWEAK in organoids increased the expression of cell proliferation and fusion markers, promoting syncytiotrophoblast formation. The source of TWEAK was found to be macrophages derived from maternal blood monocytes that adhere to sites of syncytiotrophoblast damage *in vivo*. This *in vitro* model that allows the exploration of syncytial biology has relevance for important pregnancy disorders including pre-eclampsia, where breaks occur more often, and vertical infections, where maternal monocytes could be a source of transmission.

## Introduction

The mammalian placenta is a transient, extraembryonic organ essential for fetal development *in utero*. Placental structure and function vary widely in different species in terms of its anatomical features and depth of invasion into maternal uterine tissue, reflecting evolutionary adaptations in maternal-fetal interactions. Human placentation is distinguished by the complex branching villous trees and the deep invasion of the extravillous trophoblast (EVT) into the uterus, resulting in direct contact between trophoblast and maternal tissues and blood (Fig. 1A) [1]. Central to the function of the villous placenta is the syncytiotrophoblast (syncytium, SYN), a terminally differentiated multinucleated cell layer generated through the continuous fusion of underlying villous cytotrophoblast (VCT). The SYN is in direct contact with maternal blood in the intervillous space and is the major site for rapid nutrient and gas exchange, and production of pregnancy hormones including human chorionic gonadotrophin (hCG) and progesterone. It is also an immune barrier to allorecognition because it lacks expression of any Human Leukocyte Antigen (HLA) molecules, the dominant ligands for T and Natural Killer cells [2–4]. The integrity and function of the SYN are vital for healthy pregnancy outcomes. Loss of SYN integrity is a pathological feature seen across the major pregnancy complications including miscarriage, pre-eclampsia, fetal growth restriction and stillbirth [5–8]. Furthermore, in Trisomy 21, fusion is impaired, underscoring the importance of understanding the mechanisms that regulate SYN formation and maintenance for ensuring healthy placental function [9, 10].

**Figure 1.**
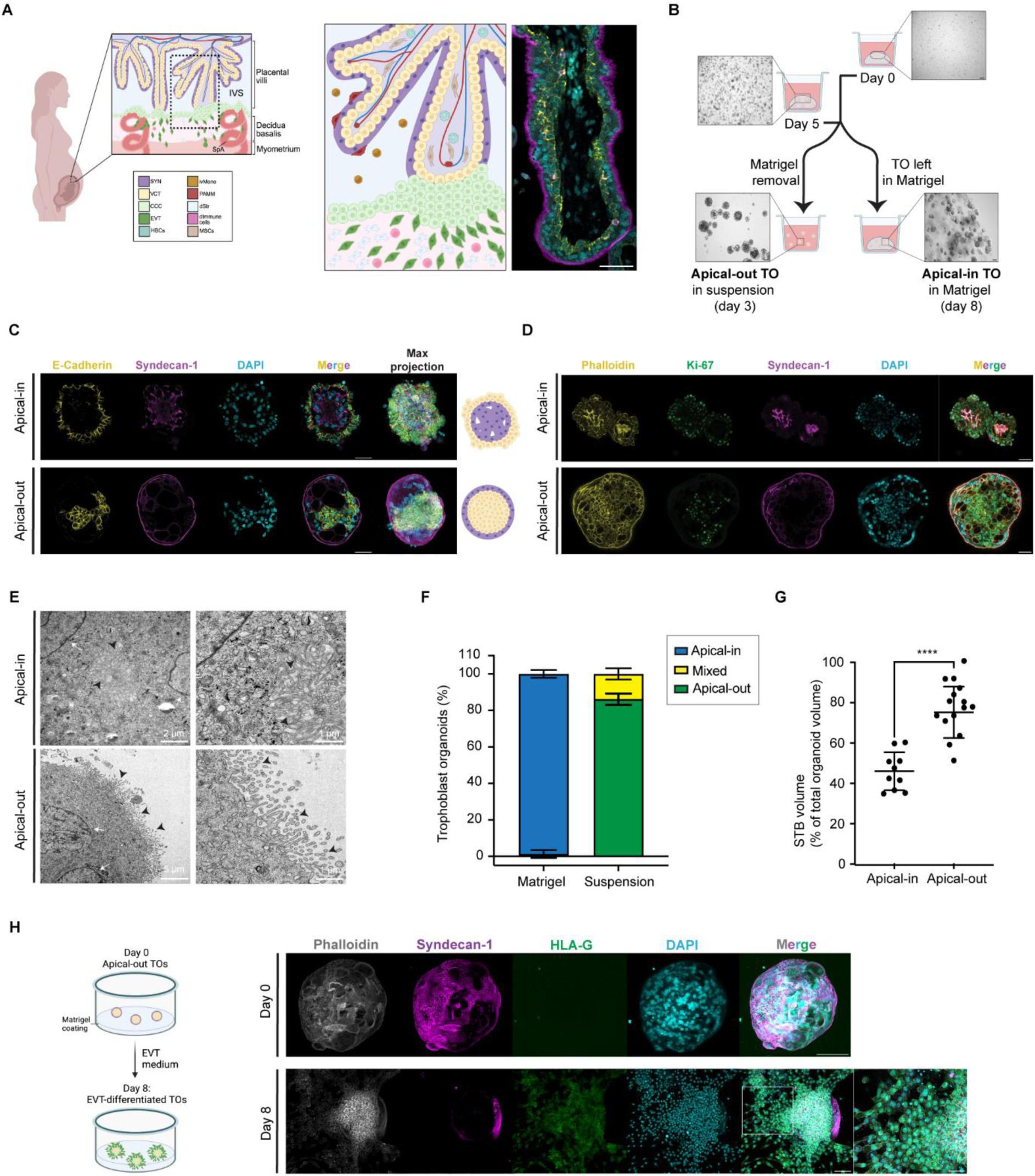
A: On the left, a schematic drawing of the maternal-fetal interface in the first trimester of pregnancy, with the major cell types present. In the zoom-in box is the structure of the placental villus, which is composed of a multinucleated outer layer, the syncytiotrophoblast (SYN), and a monolayer of villous cytotrophoblast (VCT) in contact with the mesenchymal core, where Hofbauer cells (HBCs), Mesenchymal stromal cells (MSCs) and fetal capillaries are localized. The villus is attached to the decidua where from the villous tips cytotrophoblast cell columns (CCC) give rise to invasive extravillous trophoblasts (EVT) which mingle with maternal immune (dImmune cells) and stromal cells (dStr) and remodel the spinal arteries (SpA). On the right, microscopy images of first trimester placental villus stained for E-cadherin, syndecan-1 and DAPI. Scale bars, 50 μm. PAMM, Placental associated maternal macrophages, ivMono, circulating monocytes in the intervillous space, IVS: intervillous space. B: Scheme describing the generation of apical-out trophoblast organoids with representative bright-field images. Scale bar, 100 µm. C: Confocal microscopy images of apical-in and apical-out trophoblast organoid stained for E-cadherin, syndecan-1 and DAPI, with merged images and max projection (representative image from *n* = 10 and *n* = 15 respectively). On the right, schematic drawing of the apical-in and apical-out organoids. Scale bars, 50 μm. D: Confocal microscopy images of apical-in and apical-out trophoblast organoid stained for Phalloidin, Ki-67, syndecan-1 and DAPI with merged images (representative image from *n* = 9 and *n* = 10 respectively). Scale bars, 50 μm. E: Transmission electron microscopy images of apical-in (centre of organoid) and apical-out (outside of organoid) trophoblast organoids. Microvilli are visible either on the centre of the organoids or on the surface of the organoids (arrowheads) and multiple nuclei are detected in the SYN (white arrows) (representative image from *n* = 6 and *n* = 12 respectively). F: Trophoblast organoids were imaged and analyzed using confocal microscopy and quantified for percentage of apical-in, apical-out, or mixed polarity both in Matrigel and suspension cultures (*n =* 87 and *n =* 83 respectively). Bars represent mean ± SD. G: Analysis of SYN volume of apical-in and apical-out trophoblast organoids *(n* = 10 and *n* = 15 respectively). Bars represent mean ± SD. The P value was determined using unpaired t test. **** p < 0.0001. H: On the left, schematic drawing of EVT differentiation culture of apical-out organoids. On the right, confocal microscopy images of apical-out trophoblast organoid in EVT medium at day 0 (on the left) and at day 8 (on the right) stained for Phalloidin, syndecan-1, HLA-G and DAPI (max projection is shown and representative image from *n* = 14 and *n* = 11 organoids respectively). Scale bars are 100 μm except magnified boxed areas with scale bar is 50 μm. A-G: Data from three independent organoid lines (X113, X087, Y031). H: Data from three independent organoid lines (X113, Y060, Z143). See also Figures S1 and Movies S1-2.

Research using primary trophoblast cells, explants isolated from first trimester and term placentas, and established cell lines, has identified the drivers of VCT fusion to form SYN, including hCG, glial cells missing 1 (GCM1), endogenous retroviral envelope proteins (ERVW-1, ERVFRD-1) and the cyclic AMP/PKA signalling cascade [11–16]. This process includes loss of proliferative activity, followed by cytoskeletal reorganization, and assembly of the fusion machinery resulting in cell-cell fusion [17, 18]. During pregnancy, syncytial damage, tears and shedding always occur. These are exacerbated when there is oxidative stress, as seen in early miscarriage or from turbulent flow in the intervillous space in pre-eclampsia due to defective remodelling of the spiral arteries [19, 20]. Most studies have focussed on the initiation of fusion. However, the mechanisms by which underlying VCT and SYN coordinate regeneration, remain poorly understood.

To interrogate the dynamic processes of the response of VCT to damage and syncytial regeneration, a model is needed that accurately recapitulates the native villous architecture and apical-basal polarity, and that can sustain continuous cycles of differentiation and renewal to permit repeated analysis of repair mechanisms. The conventional way of culturing trophoblast organoids (TOs) derived from first trimester placental tissue uses extracellular matrix (ECM) (for e.g. Matrigel) to provide a 3D environment; this results in the formation of the SYN internally due to the basal side of VCT interacting with the ECM [21]. Therefore, from the structural point of view, the TOs exhibit reversed polarity compared to the *in vivo* villus (hereafter apical-in). This limits experimental access to the syncytial surface, constraining mechanistic and functional analyses. Whilst recent advances, including engineered matrices and novel culture approaches including suspension culture, have begun to mitigate these constraints for several tissues including intestine, liver, and term placenta, first-trimester TOs with a more physiological configuration (apical-out) have not yet been established [22–24]. Thus there is a need for improved protocols to reproducibly generate trophoblast models capable of supporting functional studies of syncytial repair in 3D [23].

To address the fundamental question of how the SYN regenerates upon damage, we have developed a physiologically relevant, apical-out trophoblast organoid culture system that faithfully recapitulates the *in vivo* villous architecture with a functional syncytial surface covering the underlying VCT. We rigorously validated our apical-out organoids using transcriptomic and microscopic analyses as well as benchmarking them to *in vivo* first trimester tissue samples. Our system enables real-time analysis of SYN regeneration and retains its essential features and functions including marker expression, formation of the microvillous surface and hormone production. Upon induction of controlled syncytial damage, we observed molecular and structural signs of regeneration. Additionally, we show that maternal peripheral blood monocytes migrate in response to damage to the villous trophoblast. This apical-out TO model will now allow investigation of SYN function in normal and disordered pregnancies [25].

## Results

### Generation of an apical-out trophoblast organoid model that retains architectural and functional features of the first trimester placenta

To investigate how the SYN regenerates after damage, we first established an apical-out TO model of first trimester placental tissue. Established apical-in TOs were cultured in Matrigel for five days after passaging in Trophoblast Organoid Medium (TOM). Matrigel was removed and TOs were transferred to low-attachment plates for three days of suspension culture (Fig. 1B). Apical-out TOs were distinguishable under bright field microscopy by the presence of lacunae-like structures on the surface, reminiscent of SYN *in vivo*, which were absent on the surface of apical-in TOs (Fig. 1B). Apical-in TOs had E-cadherin-positive VCT on the outer surface and Syndecan-1 (SDC-1) positive SYN on the inside (Fig. 1C and Movie S1). In contrast, apical-out TOs displayed reversed polarity (Fig. 1C and Movie S2). Apical-out TOs retained proliferative VCT inside, as shown by Ki-67 staining in the core of the TOs (Fig. 1D), and SYN expressed hCGß on their outer surface (Fig. S1A). Transmission Electron Microscopy (TEM) further revealed a continuous multinucleated SYN layer with microvilli on the surface, facing the culture media, whereas apical-in TOs showed microvilli on the inside of the organoids (Fig. 1E). To validate the efficiency of our protocol, we performed 3D reconstruction of TOs in Matrigel or suspension culture using confocal microscopy and counted the number of TOs based on their polarity (apical-in, apical-out or mixed) and measured the SYN volume as a percentage of the total organoid volume. We found that 86.2% of TOs in suspension culture showed an apical-out polarity and an increased SYN volume compared to apical-in organoids (Fig. 1F-G). Together, these data demonstrate that suspension culture efficiently generates apical-out TOs, exposing a healthy syncytial surface for direct analysis.

In addition to SYN, human trophoblast also differentiates into the invasive extravillous trophoblast (EVT), which anchors the villi to the uterus *in vivo* and invades decidua to transform the maternal spiral arteries (Fig. 1A). Protocols to generate EVT from apical-in TOs are available [21, 26]. Apical-out TOs, when re-embedded in Matrigel, reverse to an apical-in configuration (Fig. S1B). Therefore, to preserve the apical-out polarity and recapitulate the *in vivo* placental villous architecture, we cultured them on a Matrigel-coated surface (Fig. 1H). After two days in trophoblast organoid medium (TOM) to sustain proliferation, apical-out TOs were exposed to extravillous trophoblast differentiation medium (EVTM) for 8 days. At day 0 in EVTM, immunofluorescence confirmed SDC-1 positive SYN on the outer surface, with no detectable HLA-G, the definitive marker of EVT. By day 8, HLA-G-positive cells migrated outwards from the apical-out TOs (Fig. 1H and S1C), demonstrating that EVT differentiation capacity is maintained while recapitulating the normal polarity of the *in vivo* placental villus structure.

We next performed bulk RNA-sequencing to characterize the time course of development of the apical-out TOs by comparing apical-out TOs after they have been transferred into suspension culture after 5 days in Matrigel (suspension day 1 to day 3) and the corresponding apical-in control in Matrigel (day 5 to day 8) (Fig. 2A). Principal component analysis (PCA) showed no major global transcriptional differences across conditions or time points, indicating preservation of the identities of the major trophoblast lineages (Fig. 2B). Core VCT markers *EPCAM* and *CDH1*, remained stable in apical-in vs apical-out TOs, while expression of SYN markers, including *ERVW-1*, *SDC1*, *PSG3* and *CGB3* were increased after three days in suspension (Fig. S1D). Quantitative real-time PCR (qPCR) analysis confirmed these findings, showing constant *MKI67* expression and progressive induction of SYN markers over time in the apical-out TOs (Fig. 2C) consistent with volumetric expansion of the SYN compartment (Fig. 1G). Analysis of differentially expressed genes comparing apical-in TOs at day 8 vs apical-out TOs at day 3 revealed upregulation of ECM-related genes (like *MMP12*, *MMP2*) and downregulation of SYN- and cytokines-related genes (like *CYP1A1*, *CCL5*) in apical-in TOs (upregulated genes in apical-in: 16, downregulated genes in apical-in: 25) (Fig. 2D). Gene ontology (GO) enrichment pointed to enrichment of processes such as estrogen metabolism and regulation of monocyte chemotaxis as enriched in apical-out TOs. Thus, in apical-out TOs, the SYN recapitulates the in vivo situation better than apical-in TOs both morphologically and functionally (Fig. 2E, Table S1).

**Figure 2.**
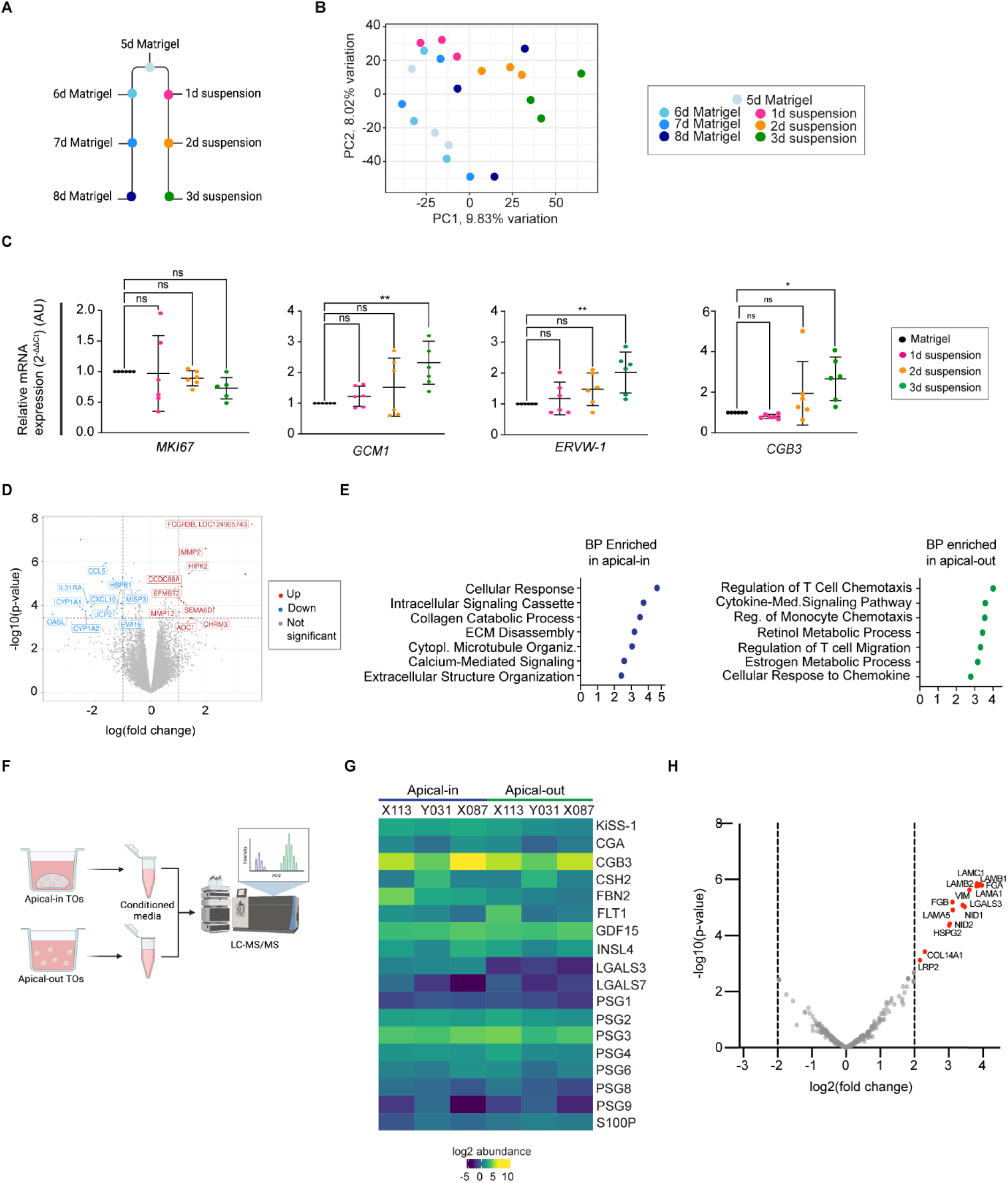
A: Scheme describing the timepoints collected for bulk RNA sequencing analysis. B: PCA of apical-in and apical-out samples analysed by bulk RNA sequencing and colour-coded based on timepoints. C: qPCR showing mRNA expression (2^-ΔΔCt^, suspension mRNA expression normalized to the corresponding control in Matrigel over the timecourse) of VCT (*MKI67*) and VCT-fusing and SYN (*GCM1, ERVW-1, CGB3)* markers in apical-in (Matrigel) and apical-out trophoblast organoids. Mean values ± SD measured in duplicates (normalized to housekeeping genes, *TBP, TOP1* and *HPRT1*) are shown. The P values were determined using one-way ANOVA multiple comparison test. ns, not significant; * p < 0.05; ** p < 0.01. AU, arbitrary units. Data from three independent organoid lines (X113, Y031, and X087). D: Volcano plot highlighting selected DEGs from the comparison between apical-in and apical-out organoids (upregulated genes: 16, downregulated genes: 25, not significant: 15699; abs(logFC) > 1, logCPM > 1, FDR < 0.05) from three independent donors (X113, Y031, and X087). E: Biological processes enriched in apical-in (on the top) and apical-out (on the bottom) trophoblast organoids. F: Scheme describing the collection of conditioned media from apical-in (day 8 in Matrigel) and apical-out (day 3 in suspension) TOs for LC-MS/MS analysis. G: Heatmap depicting the log2 abundance of selected hormones of conditioned medium from apical-in and apical-out trophoblast organoids analyzed by LC-MS/MS from three independent donors (X113, Y031, and X087). H: Volcano plot depicting enriched proteins (red dots) in apical-in vs apical-out trophoblast organoids in conditioned medium and analyzed by LC-MS/MS. Adj.pvalue threshold = 0.02, |log2FC| threshold ≥ 2. See also Figures S1 and Tables S1-S2.

The SYN is an endocrine tissue, producing steroids and peptides hormones essential for orchestrating maternal adaptations to pregnancy [27]. To assess whether apical-out TOs retain this activity, we profiled their secretome by proteomic analysis. Conditioned media from apical-in (Matrigel day 8) and apical-out (Matrigel day 5 + suspension culture day 3) cultures were analysed by liquid chromatography–tandem mass spectrometry (LC–MS/MS) (Fig. 2F). We identified 862 proteins among which pregnancy-related peptides, including Pregnancy-Specific Glycoproteins PSG2 and PSG3, Growth and Differentiation Factor 15 GDF15, Early Placental Insulin-like protein INSL4, and hCG subunits (like CGA and CGB3) (Fig. 2G, Table S2). We confirmed hCG production by ELISA in both apical-in and apical-out TOs (Fig. S1E). As expected, in apical-in TOs we found ECM-related peptides (e.g. LAMC1, NID, VIM) due to the presence of Matrigel (Fig. 2H), which were absent in apical-out TOs. These findings demonstrate that apical-out TOs preserve SYN endocrine function while avoiding confounders derived from the ECM.

In summary, we established a reproducible and efficient protocol to generate apical-out TOs from established TO lines derived from first trimester placental tissues. These retain proliferative VCT internally while fusing to form a functional SYN surface on the outside. These organoids retain lineage identity, secrete pregnancy hormones and differentiate into EVT while maintaining a villous-like organization.

### SYN fully regenerates upon induced injury in apical-out trophoblast organoids

Beyond its endocrine role, the SYN is the primary barrier protecting the fetus from any pathogens and xenobiotics present in the maternal blood. This barrier is continuously maintained during pregnancy, and damaged or aged syncytia are shed into the maternal circulation to maintain homeostasis [28]. Regeneration of the breaches is needed to prevent pathogens crossing into the feto-placental unit and compromise nutrient, gas and waste exchange [19, 29, 30]. Due to the lack of suitable *in vitro* models, this process has remained largely unexplored.

To address this, we established a protocol to mimic SYN damage using our apical-out TOs. Because high-velocity maternal blood flow arising from deficiently remodelled spiral arteries exerts elevated shear stress on the villous surface and leads to syncytial tears and microlesions *in vivo*, we employed mechanical disruption as a physiologically relevant means to model SYN damage *in vitro* [31]. We evaluated SYN regeneration in apical-out TOs at several timepoints up to 72 hours after damage (Fig. 3A). SDC-1 marking SYN, is present across the entire surface of TOs before damage, lost upon disruption, and progressively restored with initial expression of SDC-1 at 24h with full restoration of the organoid surface by 72h (Fig. 3B). Mechanical damage could be repeated multiple times (up to 3 consecutive rounds), with complete regeneration of the SYN (Fig S2A). Ultrastructural analyses using Scanning Electron Microscopy (SEM) and Transmission Electron Microscopy (TEM) confirmed this process: the surface of organoids before damage has continuous microvilli which are lost upon damage (Fig. 3C and S2B). Microvilli gradually reformed as early as 24 hours after damage, emerging beneath the degenerating outer trophoblast layer consistent with *in vivo* observations [28, 32]. By 72 hours, the surface is fully covered by a continuous, fully regenerated SYN with microvilli (Fig. 3C and S2B). hCG secretion is lost after damage but is restored after 72h showing there is also functional recovery (Fig. 3D).

**Figure 3.**
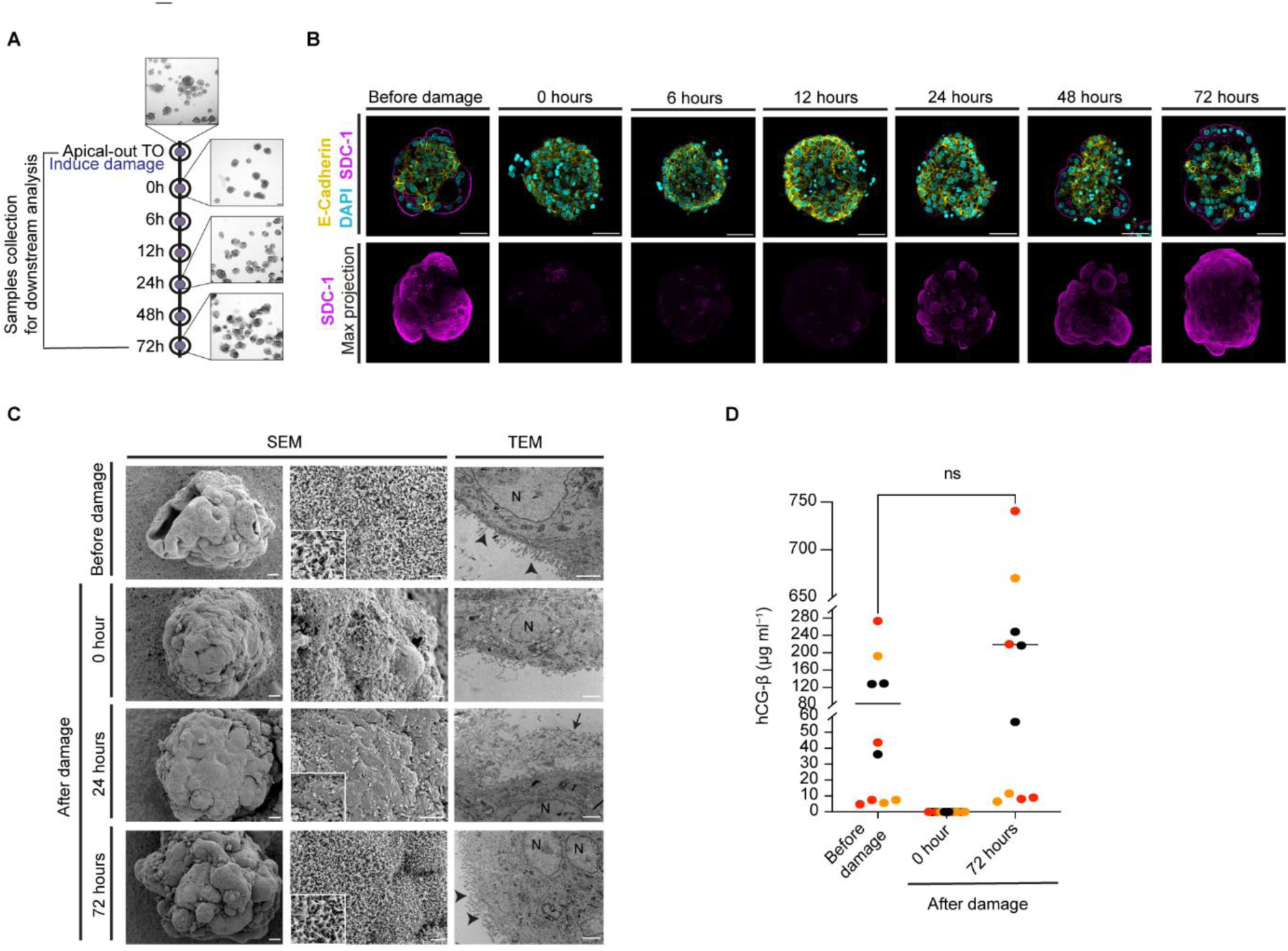
A: Scheme and representative bright-field images of apical-out trophoblast organoids before damage and after damage (0h, 24h and 72h) to mimic SYN shedding. Scale bar 100 µm. B: Confocal microscopy images of apical-out trophoblast organoid time-course before and after damage (0h, 6, 12h, 24h, and 72h) stained for E-cadherin, syndecan-1 (SDC-1) and DAPI; on the right panel, maximum projection for syndecan-1 (SDC-1) in apical-out trophoblast organoid before and after damage (0h, 6h, 12h, 24h, and 72h). Data from three independent donors (Z143, X113, Y060) Scale bars, 50 μm. C: Scanning and Transmission electron microscopy (SEM and TEM respectively) time-course images of apical-out trophoblast organoids before damage and after damage (0h, 6, 12h, 24h, and 72h). Arrowheads point to microvilli on the surface of organoids, arrow to the outer trophoblast layer shedding and each “N” marks a nucleus. Data from two independent donors (X113, Y060). Scale bars in the first column are 10 µm, and in the second and third columns they are 2 µm. D: hCG-β ELISA from conditioned medium of apical-out trophoblast organoids before damage and after damage (time point 0h and 72h). Mean values are represented and color coded by donors. Data are normalized to RNA concentration. The P value was determined using paired t test between before damage and after damage at 72h. ns, not significant. Data from three independent donors (X113, Y060, Z143). See also Figures S2.

Together, these findings demonstrate that apical-out TOs can model SYN damage and regeneration at both morphological and functional levels.

### Early transcriptional response to syncytial damage involves stress signalling, tissue remodelling and immune recruitment

To dissect the dynamics of SYN regeneration after injury, we performed bulk RNA sequencing of the intact apical-out TOs (referred to as ‘before damage’) and after mechanical disruption (0h, 6h, 12h, 24h, 48h and 72h) (Fig. 3A). PCA showed that early post-damage timepoints (6h and 12h) are clustering closely together (Fig. 4A). Differential gene expression analysis across timepoints shows that canonical SYN markers (*ERVW-1*, *CSH-1*, *SDC-1*, *CGB3*, *CYP19A1*, *CGA)* are highly expressed in intact organoids, lost after damage and progressively re-expressed starting from 24h and returning to baseline by 72h highlighting a clear regenerative trajectory (Fig. 4B-C, S3A, and Table S3). This protocol captures the temporal induction of known SYN regulators including *GCM1* as well as novel candidates such as *EPS8L1*, which lies within the chromosome 19q13.4 region that harbours a cluster of genes associated with reproduction (Fig. 4B, S4A and Table S3) [11, 33]. EPS8L1 expression was confirmed at the protein level to be specifically expressed in SYN, by immunohistochemistry (IHC) in organoids and first trimester placental tissues (Fig. S4C).

**Figure 4.**
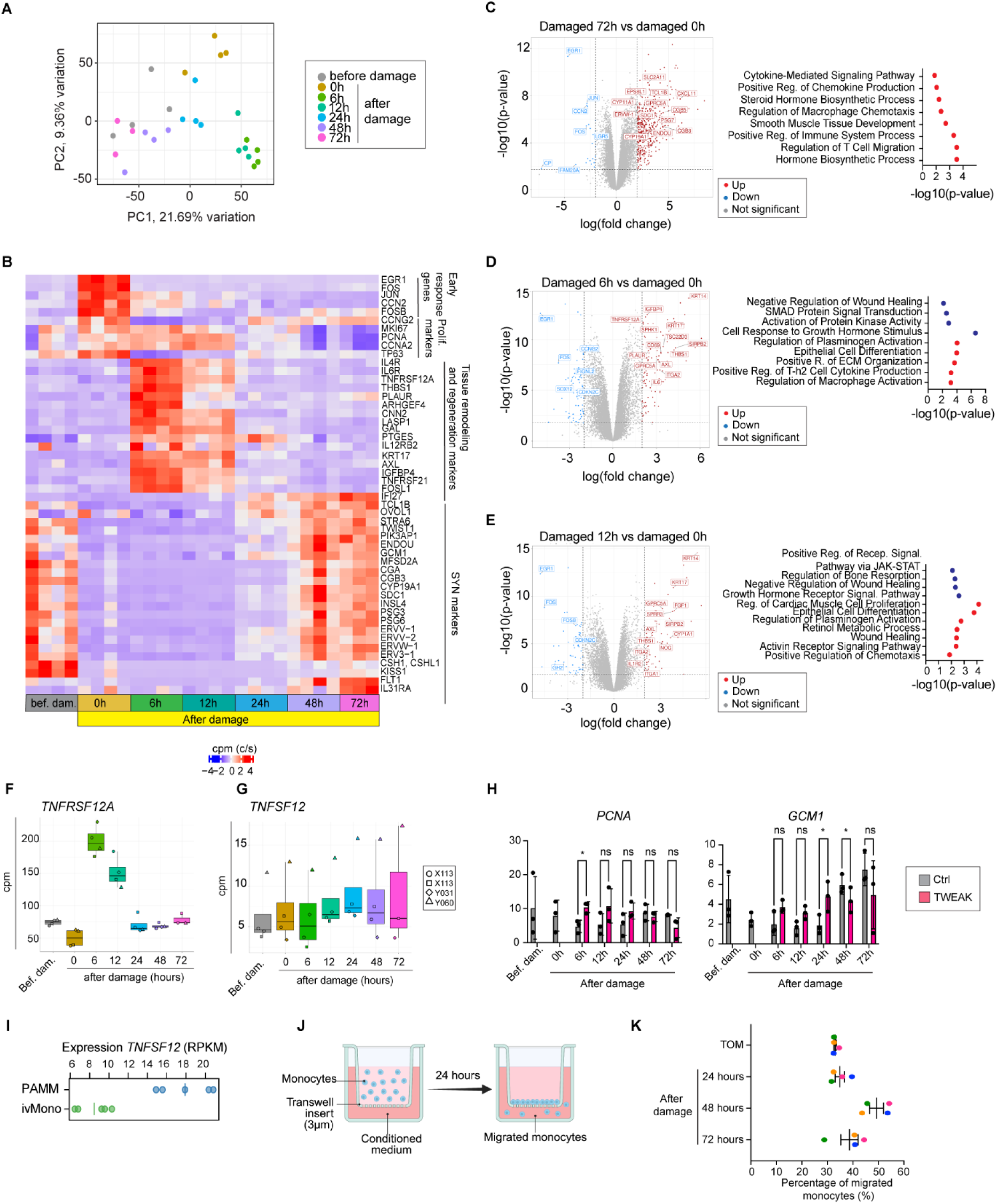
A: PCA of apical-out trophoblast organoids samples analysed by bulk RNA sequencing where colours indicate time-points. Data from three independent donors (X113, Y060, Y031). Experiment with donor X113 has been repeated twice. B: Heatmap depicting centered and scaled counts per million (cpm) of selected genes expressed in apical-out trophoblast organoids before and after damage in the time-course experiment. C: On the left, volcano plot highlighting selected DEGs from the comparison between damaged organoids at time point 72h vs damaged organoids at 0h (upregulated genes: 533, downregulated genes: 25, not significant: 15146; abs(logFC) > 2, logCPM > 1, FDR < 0.05). On the right, biological processes upregulated in damaged organoids at time point 72h after damage. D: On the left, volcano plot highlighting selected DEGs from the comparison between damaged organoids at time point 6h vs 0h (upregulated genes: 114, downregulated genes: 89, not significant: 15501; abs(logFC) > 2, logCPM > 1, FDR < 0.05). On the right, biological processes downregulated (in blue) and upregulated (in red) in damaged organoids at time point 6h. E: On the left, volcano plot highlighting selected DEGs from the comparison between damaged organoids at time point 12h vs 0h (upregulated genes: 78, downregulated genes: 72, not significant: 15554; abs(logFC) > 2, logCPM > 1, FDR < 0.05). On the right, biological processes downregulated (in blue) and upregulated (in red) in damaged organoids at time point 12h. F: Expression level (in cpm) of *TNFRSF12A* gene across the time-course experiment. G: Expression level (in cpm) of *TNFSF12* (*TWEAK*) gene across the time-course experiment. H: Expression level of TWEAK (TWEAK) in PAMM and ivMono. Each dot represents an individual donor (n=5), and vertical lines indicate the mean across donors. I: qPCR showing mRNA expression of *PCNA* and *GCM1* in control and TWEAK-treated TOs. Mean values ± SD measured in duplicates (normalized to housekeeping genes, *TBP, TOP1* and *HPRT1*) are shown. The P values were determined using multiple comparison test. ns, not significant; * p < 0.05. Data from three independent experiments. J: scheme describing the CD14+ monocytes migration assay using transwell and conditioned medium from damaged organoids. K: Percentage of CD14+ monocytes that migrated in the transwell assay. Mean values and SEM are represented. Each color indicates an independent experiment. See also Figures S3-S5 and Tables S3.

In contrast to SYN markers, proliferation markers (*MKI67*, *PCNA*) are highly expressed between 0-12h and declined as VCT fusion is initiated from 24h, reflecting the proliferation of the remaining VCT and the absence of SYN (Fig. 4B). Immediate-early response genes (*EGR1*, *FOS*, *JUN*, *FOSB)* are rapidly induced at 0h with an enrichment of genes involved in biological processes associated with stress response signalling and regulation of macrophage chemotaxis (Fig. 4B and S3B). This signature is attenuated by 6-12h, during which there is a shift in expression of genes involved in tissue remodelling and wound healing (*THBS1, PLAUR*) (Fig. 4B, 4D-E and S4B). We observed expression of thrombospondin 1 (*THBS1*) at 6h at 12h after damage, while no signal was detectable before damage and from 24h after damage (Fig. S4D). This was validated at the protein level by IHC in the TOs over the timecourse experiment and first trimester placental tissue where we see expression of THBS1 in villous stroma and in VCT at sites of damage (Fig. S4D).

Several cytokine receptors (*IL4R*, *IL6R*, *TNFRSF12A*, *TNFRSF21*) are also upregulated during the regenerative window, suggesting activation of immune recruitment programs at the damaged surface (Fig. 4B, 4D-F, and S5A). The expression of these receptors then decreases to a level similar to the ‘before damage’ condition, suggesting their expression is triggered specifically by the damage (Fig. 4F and S5A).

Epithelial differentiation and tissue remodelling processes were significantly enriched in damaged organoids at 24-48h compared with those at 0h, indicating response to the injury and differentiation of VCT to reform SYN (Fig. S3C-D). By 72h, damaged and intact control organoids revealed similar transcriptomes with only a small number of upregulated genes (94/15704), including *IL2RG*, *IL31RA*, *OAS1*, *IFI27*, *IFITM1*, which are involved in the type I Interferon-mediated signalling pathway and the antiviral innate immune response (Fig. S3A and Table S3).

In summary, we show that apical-out TOs mount a coordinated transcriptional program of proliferation and differentiation towards the SYN with full barrier restoration by 72h providing a robust model to investigate mechanisms of SYN regeneration.

### TWEAK supplementation enhances SYN regeneration

Next, we aimed to identify how SYN regeneration is regulated. Among the genes upregulated in the early window after injury, *TNFRSF12A* emerged as a candidate regulator of syncytial regeneration (Fig. 4F). *TNFRSF12A* is induced by injury, oxidative stress and inflammation [34]. Its activation upon binding to TNFSF12 (also known as TWEAK) engages multiple intracellular signalling pathways, including nuclear factor-κB (NF-κB) [35]. TWEAK itself is primarily expressed by leukocytes, including macrophages, and has been implicated in tissue repair and regeneration in muscle and liver, with no information reported in the human placenta [36–39]. This prompted us to further investigate the role of TWEAK in SYN regeneration, hypothesizing that trophoblast injury induces upregulation of TNRFSF12A to enable a response to damage and recruit maternal leukocytes. Indeed, in our bulk RNA-seq data the gene encoding for TWEAK, *TNFSF12,* is expressed at very low levels (Fig. 4G). This raised the question whether the supplementation of human recombinant TWEAK in the TOM could stimulate an earlier SYN regeneration in damaged organoids. Indeed, qPCR analysis shows that treatment with TWEAK enhanced early proliferative response with an upregulation of the S-phase marker *PCNA* as early as 6h compared to non-treated TOs (Fig. 4H). There is also a statistically significant increase in expression of *GCM1* at 24h after damage (Fig. 4H). Other genes involved in cell fusion and SYN markers like *GJA1*, *ERVW-1* and *SDC-1* show a similar increasing trend (Fig. S5A).

We also explored whether other cytokines might increase cell proliferation and enhance SYN regeneration. *IL4R* and *IL6R* genes showed increased expression at 6-12h after damage, while *IL4* and *IL6* genes encoding for their respective ligands are not expressed in our bulk RNA seq data (Fig. S5B-C). In contrast to TWEAK, supplementation with human recombinant IL-4 and IL-6 did not show a significant upregulation of cell proliferation (*MKI67, PCNA*), cell-cell fusion (*GCM1*) or SYN (*ERVW-1 or CGB3*) markers but instead we observed an opposite trend compared to the TWEAK treated TOs (Fig. S5D-E). The increase in proliferation and cell-cell fusion observed after supplementation with TWEAK suggests that it is involved in SYN regeneration after damage as seen in other tissues.

The changes in gene expression observed in damaged TOs following recombinant TWEAK treatment, prompted us to investigate the potential external sources of TWEAK (Fig. 4F). Placenta-associated maternal macrophages (PAMM, also known as PAMM1a) are known to adhere to SYN at sites of damage and do secrete factors involved in tissue repair [40]. We therefore looked for *TNFSF12* expression in the placental-associated myeloid cells and found that *TNFSF12* is expressed in PAMM, while its expression is very low in circulating monocytes in the intervillous space (ivMono, also known as PAMM1b) (Fig. 4I).

We next examined whether damaged TOs are able to attract ivMono through injury-induced upregulation of chemoattractants, as suggested by our RNA-seq data (Fig. 4D,E). To assess this, we performed a migration assay where CD14+ primary monocytes isolated from peripheral blood mononuclear cells (PBMCs) were seeded on top of a 3μm insert in the upper chamber, and conditioned media from damaged organoids was loaded in the lower chamber (Fig. 4J). An increase in monocytes migration is seen when monocytes are exposed to conditioned medium from TOs 48h after damage, suggesting damaged organoids do attract monocytes to adhere to the injured SYN (Fig. 4K).

The finding that TWEAK is expressed by PAMM and enhances cell proliferation and SYN formation, underscore the role of PAMM in SYN regeneration after damage.

## Discussion

This study describes a novel trophoblast organoid model to study how human SYN regenerates after damage. We first developed an apical-out organoid model that more accurately recapitulates the cellular composition, structure and physiology of the first trimester human placenta. Transcriptomic data showed that apical-out TOs retain the expression of canonical trophoblast markers. Furthermore, unbiased proteomic analysis shows that the apical-out TOs retain the secretory and endocrine function of the SYN, releasing multiple glycoproteins such as pregnancy-specific glycoproteins (PSGs) and hCG. Additionally, to visualize EVT differentiation, thereby recapitulating the complete VCT differentiation pathway, we established a protocol in which apical-out TOs are seeded on a thin Matrigel layer. While maintaining their apical-out configuration, EVT migration and invasion into the surrounding matrix is seen.

We next used our model to study how the SYN is regenerated following the inevitable damage that occurs throughout pregnancy due to the SYN being continuously exposed to physical and oxidative stress, leading to microlesions and shedding of syncytial fragments particular under pathological conditions such as PE [19]. To study this regenerative response *in vitro*, we applied controlled mechanical disruption to apical-out TOs and monitored their recovery over 72 hours. Signs of regeneration were evident by 24h, with a gradual upregulation of cell-cell fusion and SYN-associated genes and complete restoration of SYN and function by 72h confirmed by electron microscopy and hCG secretion.

Between 6-12h following damage, we detected increased expression of cytokines receptors like *IL4R*, *IL6R*, and *TNFRSF12A.* Because TOs do not express the corresponding ligands, we tested whether exogenous supplementation could enhance SYN regeneration. While IL-4 and IL-6 supplementation had no significant effect on proliferation, fusion, or SYN markers expression, addition of TWEAK, the ligand of TNFRSF12A, significantly increased *PCNA* expression, suggesting enhanced DNA replication and cell proliferation, as well as expression of the fusion marker *GCM1*. These findings are consistent with studies in other system where TWEAK has been showed to enhance muscle and liver regeneration [37–39]. As TWEAK was not expressed in our TOs we showed that PAMM, previously shown to adhere on damaged SYN *in vivo*, are the source of TWEAK and are involved in SYN regeneration. We therefore established a migration assay using transwells and medium from damaged organoids, and we showed that CD14+ primary monocytes respond to migration cues present in the conditioned media.

Our apical-out TO model provides a valuable tool to study the function of the SYN in a range of conditions that are important in pregnancy. For example, because the placenta experiences fluctuating oxygen levels *in vivo*, the direct exposure of the SYN to the culture medium allows investigation of how oxygen regulates of trophoblast differentiation. The model could also be used to study transmission of pathogens across the placenta. Because most pathogens that cause vertical infections are present inside macrophages (for e.g. CMV, Toxoplasma, Listeria) the maternal PAMM that adhere to damaged SYN could be entry routes into the fetus [41]. Culturing infected primary monocytes with our damaged TOs could help understand whether they are the vehicle for viral transmission into the placenta.

Overall, this work describes an ill-understood area of SYN biology. Understanding of the response to inevitable SYN damage that always occurs during any pregnancy and the role of PAMM and signalling factors like TWEAK that contribute to this process, has implications for many pregnancy disorders, particularly vertical transmission and pre-eclampsia. These first trimester apical-out TOs could also be used in toxicology studies for novel therapeutics, as well as in the safety evaluation of therapies for pregnant women with chronic conditions, such as diabetes and obesity.

## Materials and Methods

### KEY RESOURCES TABLE

| REAGENT or RESOURCE | SOURCE | IDENTIFIER |
| --- | --- | --- |
| **Antibodies** |  |  |
| Anti-SDC1 (44F9, mouse) | Miltenyi Biotec | Cat# 130-108-028;<br>RRID:AB_2655047 |
| Anti-SDC1 (E7F7T, rabbit) | Cell Signalling | Cat# 96489;<br>RRID:AB_3713648 |
| Anti-E-Cadherin (24E10, rabbit) | Cell Signalling | Cat# 3195;<br>RRID:AB_2291471 |
| Anti-Ki67 (SP6, rabbit) | Invitrogen | Cat# MA5 – 14520;<br>RRID:AB_10979488 |
| HLA-G antibody (G233, mouse) A488 | Novusbio | Cat#<br>NB110-55298AF488;<br>RRID:AB_3177325 |
| Anti-hCGb (5H4-E2, mouse) | Abcam | Cat# Ab9582;<br>RRID:AB_296507 |
| THBS1 (rabbit, R06-5F3) | Abnova | Cat# RAB01344<br>RRID:AB_3709914 |
| TWEAKR/Fn14 (rabbit, polyclonal) | Cell Signalling | Cat# 4403<br>RRID:AB_10693941 |
| EPS8L1 (rabbit, polyclonal) | Atlas Antibodies | Cat# HPA041851<br>RRID:AB_2677706 |
| Secondary ab- Donkey anti-mouse 488 | Invitrogen | Cat# A32766;<br>RRID:AB_2762823 |
| Secondary ab- Donkey anti-mouse 568 | Invitrogen | Cat# A10037;<br>RRID:AB_11180865 |
| Secondary ab- Donkey anti-rabbit 568 | Invitrogen | Cat# A10042;<br>RRID:AB_2534017 |
| Secondary ab- Donkey anti-mouse 594 | Invitrogen | Cat# A21203;<br>RRID:AB_2535789 |
| Peroxidase AffiniPure® F(ab')<sub>2</sub> Fragment Donkey Anti-Rabbit IgG (H+L) | Jackson ImmunoResearch Laboratories, Inc. | Cat# 711-036-152 |
| **Biological sample: primary trophoblast organoids and primary monocytes** |  |  |
| Primary trophoblast organoids (TOs); female | Addenbrooke's Hospital | X113 |
| Primary trophoblast organoids (TOs); male | Addenbrooke's Hospital | Z143 |
| Primary trophoblast organoids (TOs); female | Addenbrooke's Hospital | Y060 |
| Primary trophoblast organoids (TOs); female | Addenbrooke's Hospital | Y031 |
| Primary trophoblast organoids (TOs); male | Addenbrooke's Hospital | X087 |
| **Chemicals, peptides, and recombinant proteins** |  |  |
| Phalloidin-647 | Cell Signalling | Cat# 8940 |
| Advanced DMEM/F12 | Life Technologies | Cat# 12634010 |
| N2 Supplement | Life Technologies | Cat# 17502048 |
| B27 Supplement minus vitamin A | Life Technologies | Cat# 12587010 |
| Primocin | Invivogen | Cat# ant-pm-1 |
| N-Acetyl-L-Cysteine | Sigma-Aldrich | Cat# A9165 |
| L-Glutamine | Life Technologies | Cat# 25030-024 |
| A83-01 | Tocris | Cat# 2939 |
| CHIR99021 | Tocris | Cat# 4423 |
| Y-27632 | Millipore Sigma | Cat# 688000 |
| rhEGF | Peprtech | Cat# AF-100-15 |
| rhHGF | Peprtech | Cat# 100-39 |
| rhR-Spondin1 | BioTechne | Cat# 4645-RS/CF |
| rhFGF2 | Peprtech | Cat# 100-18B |
| Prostaglandin E2 | Millipore Sigma | Cat# P0409 |
| RhTWEAK | Peprtech | Cat# 310-06 |
| rhIL-6 | Peprtech | Cat# 206-IL |
| rhIL-4 | Peprtech | Cat# 204-IL |
| 2-mercaptoethanol | Gibco | Cat# 51500-056 |
| Bovine serum albumin | Sigma-Aldrich | Cat# A8412 |
| ITS-X supplement | Gibco | Cat# 51500-056 |
| Knockout serum replacement | Thermo Fisher | Cat# 10828028 |
| Hi FBS | Gibco | Cat# 12634010 |
| Amphotericin B | Sigma-Aldrich | Cat# A9528 |
| Gentamycin | Sigma-Aldrich | Cat# G1272 |
| HEPES | Gibco | Cat# 15630106 |
| M-CSF | Peprtech | Cat# 300-25 |
| Accutase | Gibco | Cat# A11105-01 |
| Cell Recovery Solution | Corning | Cat# 354253 |
| Paraformaldehyde | Thermo Fisher | Cat# 47347.9M |
| Poly-L-lysine | Gibco | Cat# A3890401 |
| Sodium deoxycholate | Sigma-Aldrich | Cat# D6750 |
| Trypsin, TPCK Treated | Thermo Fisher Scientific | Cat# 20233 |
| Lysyl Endopeptidase®, Mass Spectrometry Grade (Lys-C) | FUJIFILM Wako Pure Chemical Corporation | Cat# 129-02541 |
| Histogel | Eprelia | Cat# HG-4000-012 |
| 10% neutral buffered formalin solution | Sigma-Aldrich | Cat# HT5011 |
| **Critical commercial assays** |  |  |
| Human hCG-beta ELISA Kit | Thermo Fisher | Cat# EH235RB |
| RNeasy Micro Kit | Qiagen | Cat# 74004 |
| TMT10plex Isobaric Label Reagent Set | Thermo Fisher Scientific | Cat# 90309 |
| SuperScript™ VILO™ cDNA Synthesis Kit | Thermofisher | Cat# 11754250 |
| TaqMan™ Fast Advanced Master Mix | Thermo Fisher | Cat# 4444964 |
| **Deposited data** |  |  |
| Bulk RNA-seq | This paper | https://www.ebi.ac.uk/biostudies/arrayexpress/studies/E-MTAB-15647 |
| Secretome analysis | This paper | https://www.ebi.ac.uk/pride/archive/projects/PXD063812 |
| **Software and algorithms** |  |  |
| ImageJ | NIH | https://imagej.nih.gov/ij/ |
| R (Version 4.5.0) | R Project | https://www.r-project.org |
| edgeR (Version 4.6.3) [42] | Bioconductor | https://bioconductor.org/packages/edgeR |
| sva (Version 3.56.0) [43] | Bioconductor | https://bioconductor.org/packages/sva |
| ComplexHeatmap (Version 2.24.1) [44, 45] | Bioconductor | https://bioconductor.org/packages/PCAtools |
| ggplot2 (Version 4.0.0) [46] | CRAN | https://bioconductor.org/packages/limma |
| PCAtools (Version 2.20.0) [47] | Bioconductor | https://bioconductor.org/packages/edgeR |
| dplyr (Version 1.1.4) [48] | CRAN | https://cran.r-project.org/package=dplyr |
| ggrepel (Version 0.9.6) [49] | CRAN | https://cran.r-project.org/package=ggrepel |
| limma (Version 3.64.3) [50] | Bioconductor | https://bioconductor.org/packages/limma |
| Prism 10 | GraphPad | https://www.graphpad.com |
| Enrichr [51] | Ma'ayan Lab | https://maayanlab.cloud/Enrichr/ |
| **Other** |  |  |
| Source code used for analysis and generation of plots | Github | https://github.com/fmicompbio/apical-out-placental-organoids |

### RESOURCE AVAILABILITY

#### Lead contact

Requests for further information should be directed to and will be fulfilled by the lead contact Margherita Y. Turco.

#### Materials availability

Requests for resources and reagents should be directed to and will be fulfilled by the lead contact Margherita Y. Turco.

#### Data and code availability

Bulk RNA-seq data have been deposited at ArrayExpress collection in BioStudies: https://www.ebi.ac.uk/biostudies/arrayexpress/studies/E-MTAB-15647. All source code used for analysis and generation of plots, as well as additional supplementary tables, are available from GitHub: https://github.com/fmicompbio/apical-out-placental-organoids. The mass spectrometry proteomics data have been deposited to the ProteomeXchange Consortium via the PRIDE partner repository [52].

### EXPERIMENTAL MODEL AND SUBJECT DETAILS

#### Derivation of primary trophoblast organoids (TOs)

Human placental tissues used in this study were obtained from elective first-trimester pregnancy terminations with written informed consent in accordance with the Declaration of Helsinki (2000). Samples were collected at Addenbrooke’s Hospital (Cambridge, UK) under ethical approvals from the Cambridge Local Research Ethics Committee (20/LO/0115 and 04/Q0108/23). Approval (04/Q0108/23) is now incorporated into the East of England– Cambridge Central Research Ethics Committee overarching ethics permission, granted to the Centre for Trophoblast Research biobank for the ‘Biology of the Human Uterus in Pregnancy and Disease Tissue Bank’ at the University of Cambridge (17/EE/0151). The transfer and use of all the placental tissue samples together with the established trophoblast organoid cultures for this study are authorized under a Material Transfer Agreements (MTA) between the University of Cambridge and the Friedrich Miescher Institute for Biomedical Research (Agreements No. G119629 and G108313). Donor information is described in Table S4.

#### Primary monocytes

Peripheral blood was taken from healthy adult volunteers (Donor IDs: #169, and #173). All samples were obtained with written informed consent from participants under ethical approval which was obtained from the Chair of the Human Biology Research Ethics Committee (study HBREC.2022.26).

### METHOD DETAILS

#### Organoid culture

TOs were grown as previously described [21]. Briefly, organoids were grown in Matrigel (Corning, 356231) droplets supplemented with trophoblast organoid medium (TOM). [Advanced DMEM/F12, 1X N2 Supplement, 1X B27 Supplement minus vitamin A, 100 μg/mL Primocin, 1.25 mM N-Acetyl-L-Cysteine, 2 mM L-Glutamine, 500 nM A83-01, 1.5 μM CHIR99021, 5 μM Y-27632, 50 ng/mL rhEGF, 50 ng/mL rhHGF, 80 ng/mL rhR-Spondin1, 100 ng/mL rhFGF2, and 2.5 μM Prostaglandin E2]. After 7-10 days TOs were first broken apart by automatic pipetting (400 times) and then incubated with pre-warmed Accutase cell detachment solution for 5 minutes at 37°C. The digest was passed through a 40 μm nylon mesh cell strainer and the flow through was mixed with Matrigel. 25μl Matrigel domes were plated in 48-well plate (Corning, 3548) and incubated at 37°C for 20min. After, 250μl of TOM was added per each well and medium was changed every 2-3 days.

#### Apical-out TOs generation

After 5 days in Matrigel, TOs were collected and Matrigel was removed using Cell Recovery Solution (Corning, 354253) for 50 min maximum on ice. TOs were washed with cold PBS pH (7.4) (GIBCO, 10010-015) and resuspended in TOM. TOs were then plated in low attachment 96-well plate (Corning, 3474) in suspension (one 48well plate spitted in 4-5 wells of the 96-well plate) and cultured at 37°C for 3 days. Bright field images were taken using EVOS XL Core microscope (Invitrogen).

#### Differentiation of Apical-out TOs into EVT lineage

96w plates were coated with a layer of 20μl of Matrigel. To ensure a flat Matrigel layer, the 96w plate is centrifuged at 1000rcf for 1minute at 4°C. Apical-out organoids at day 3 were filtered with 100 μm nylon mesh cell strainer and organoids with size above 100 μm were plated on top of a Matrigel coated 96w plate. Organoids were cultured in TOM for two days before switching to EVTM for 8 days [Advanced DMEM/F12, 0.1mM 2-mercaptoethanol, 2 mM L-Glutamine, 100μg/mL Primocin, 0.3% bovine serum albumin, 1% ITS-X supplement, and 4% knockout serum replacement]. Medium was changed every 2 days. Bright field images were taken using Nikon Eclipse Ts 2R microscope.

#### Apical-out damage protocol

After 3 days in suspension culture, apical-out organoids were filtered with a 100 μm nylon mesh cell strainer and organoids with size above 100 μm were either collected for further analysis (referred as before damage or not damage) or subjected to the damage protocol: For damage, TOs were subjected to automatic pipetting (400X) using a small-bore pipette tip (Metttler Toledo, BioClean, RT-200FLR) and then pipetted manually 100X. TOs were passed through a 40 μm nylon mesh cell strainer to remove detached SYN and cell debris, and organoids above 40 μm were plated in 96-wells low attachment plates and incubated at 37°C for other 3 days. In the cytokines treated experiments, TOs at 0h after damage have been cultured in the presence of IL-6 (50ng/ml), IL-4 (50ng/ml), and TWEAK (100ng/ml) for 3 days. Bright field images were taken using EVOS XL Core microscope (Invitrogen).

#### Isolation of primary monocytes

Adult blood was obtained by venepuncture into EDTA collection tubes (BD). Blood samples were diluted 1:2 in phosphate buffered saline (PBS), layered onto a Pancoll gradient (PAN biotech, P04-60500) and centrifuged at 3000 rcf for 20 minutes without brake. The PBMC interface was collected and washed in PBS. Platelets were depleted via centrifuging at 200G for 10 minutes without brake. Cells were resuspended then counted using a haemocytometer and Trypan-blue (ThermoFisher, 15250061). CD14^+^ monocytes were enriched using the CD14 MicroBead kit (Miltenyi, 130-050-201) as per the manufacturer’s instructions. Briefly, cells were blocked with FACS buffer at 80μL per 10million cells and labelled with CD14 MicroBeads at 20μL per 10million cells for 15 minutes at 4°C. Cells were washed with FACS buffer to remove unbound beads and centrifuged at 400G for 5 minutes at 4°C. Cells were resuspended in 500μL of FACS per 100million cells. A suitably sized MACS separator and column were assembled. The MS column (Miltenyi, 130-042-201) was used. Columns were prepared with FACS buffer, and the cell suspension washed through. Unlabelled cells were washed through with 3 washes through the column. 1ml of FACS buffer for MS columns was added, the column quickly removed from the magnet, placed over a 15ml Falcon tube and cells plunged from column into tube. The eluted CD14^pos^ cells were counted and stored at 5-50million cells per ml in CryoStor CS10 (STEMCELL Technologies, 100-1061) in liquid nitrogen until use.

#### Primary monocytes culture

Primary monocytes were cultured in 96-wells plate at 50-100 × 10^5^ confluency in Primary Monocytes Medium (PMM) [Advanced DMEM/F12, 10% Hi-FBS, 10U/ml Pen/strep, 2.5μg/ml Amphotericin B, 0.5μg/ml Gentamycin, 10mM L-Glutamine, 10mM HEPES, and 100ng/ml M-CSF]. Medium was changed every 2 days.

#### Monocytes migration assay

Primary monocytes (50 × 10^5^ cells resuspended in 100μl) were plated on the 3μM insert (Thermo Fisher, Cat. 140629) and incubated for 10 minutes at 37°C. In the lower chamber, 600μl of conditioned medium from damaged TOs was loaded. TOM was used as a control, after 24h incubation, cells in the lower chamber were counted with the NucleoCounter NC-202 (Chemometec).

#### Illumina stranded mRNA-seq library preparation and bulk RNA-sequencing

cDNA libraries were prepared using the Illumina Stranded mRNA-Seq (Illumina IDT DNA/RNA UDI) preparation kit according to the manufacturer’s instructions. Libraries were pooled and sequenced with the Illumina NovaSeq 6000 platform, producing paired end reads of 56 base pairs each (2×56bp). Demultiplexing and FASTQ file generation were performed using Illumina’s bcl2fastq2 software.

#### Immunofluorescence

Apical-out and apical-in TOs were fixed in 4% PFA (Thermo Fisher, 47347.9M). Permeabilization has been performed using 0.5% Triton for 45min. Fixed and permeabilized TOs were then plated on plasma treated and poly-L-lysine (Gibco, A3890401) coated µ-Slide 8 Well Glass Bottom dishes (Ibidi, 80807). After 1 hour, blocking was performed using 3% Donkey Serum (Pan Biotech, P30-0101) in 1X PBS. TOs were incubated with primary antibodies (listed in Key resource table) in antibody buffer solution (1.5% Donkey Serum + 0.05% triton) overnight at 4°C. After four washes with 1X PBS, TOs were incubated with secondary antibodies (listed in key resource table) in antibody buffer solution (1.5% Donkey Serum + 0.05% triton) overnight at 4°C or 3h at RT. After four washes with 1X PBS, imaging was performed confocal microscope Stellaris 5 with a 20X objective (Leica). For SYN volume quantification, after immunostaining organoids were cleared in a Ce3D solution [(*N*-methylacetamide (Sigma-Aldrich, M26305-500G), Histodenz (Sigma-Aldrich, D-2158), 1-Thioglycerol (Sigma-Aldrich, M1753) and Triton X-100] overnight and then imaging was performed using 2-Photon Stellaris 8 microscope with a 25X objective (Leica). Antibodies dilutions are listed in Table S5.

#### Fixation and paraffin embedding of TOs

TOs were collected at different timepoints. For apical-in TOs Matrigel was removed using Cell Recovery Solution (Corning, 354253) for maximum 50 min on ice; for apical-out TOs were collected by centrifugation. TOs were washed with cold PBS pH (7.4) (Gibco, 10010-015) and fixed in 10% neutral buffered formalin solution for 45 min at RT. Fixed TOs were centrifuged at 300 rcf, washed twice in PBS and embedded in pre-warmed, liquid Histogel. After solidification at RT, the Histogel dome containing TOs was transferred into a labelled histology cassette (Cell Path, EAM-0309-72B) and stored in 70% ethanol until dehydration and paraffin embedding. TOs were loaded in the retort of the HistoCore PEARL tissue processor (Leica, 14 0493 50667) using overnight standard tissue dehydration method until paraffin inclusion. The samples were then removed from the tissue processor, transferred in molten paraffin and attached to the cassettes using the HistoCore Arcadia Embedding Center (Leica, 14 0393 57262 and 14 0393 57257). After solidification of the paraffin blocks, microtome (Epredia, HM 355S) sectioning was performed.

#### Immunohistochemistry

Immunohistochemistry was performed on 3-μm-thick paraffin sections of both placental tissues and TOs using the Leica Bond RX automated stainer. As part of the automated staining, the sections were first deparaffinized with the BOND Dewax Solution (Biosystems Switzerland AG, AR9222) and cleared with 100% ethanol. Next, heat induced epitope retrieval was performed by boiling the sections in BOND Epitope Retrieval Solution 1, pH6 (Biosystems Switzerland AG, AR9961) or BOND Epitope Retrieval Solution 2, pH9 (Biosystems Switzerland AG, AR9640) at 100°C for 10min. Sections were incubated with appropriate primary antibody dilutions for 60 minutes at room temperature followed by washes with BOND Wash Solution 1X (BOND Wash Solution 10X Concentrate, Biosystems Switzerland AG, AR9590). The binding sites of the primary antibodies were then revealed using the ready-to-use Bond Polymer Refine detection kit (Biosystems Switzerland AG, DS9800) containing peroxide block solution, post primary, polymer reagent, DAB chromogen and hematoxylin counterstain. Sections were then, dehydrated with consequent 5-minute incubations in 70% ethanol, 95% ethanol and 100% ethanol. Last, sections were dried at room temperature prior to addition of Tissue-Tek® Glas™ Mounting Medium (Sakura, 1408N). Coverslips 24×50mm #1,5 (Epredia, BB02400500SC13MNZ0) were applier using Tissue-Tek® Glas™ g2-E2 instrument (Sakura). Stained sections were imaged with the Zeiss Axioscan Z1 slide scanner. Antibodies dilutions are listed in Table S5.

#### 2-plex fluorescent immunohistochemistry

Multiplex immunofluorescence staining was performed with the TSA-based Opal fluorescence technology (Akoya Biosciences, Marlborough, MA) on 3-µm thick paraffin sections of both placental and TOs using the Leica Bond RX automated stainer. As part of the automated staining, the sections were first deparaffinized with the BOND Dewax Solution (Biosystems Switzerland AG, AR9222) and cleared with 100% ethanol. Next, heat induced epitope retrieval was performed by boiling the sections in BOND Epitope Retrieval Solution 2, pH9 (Biosystems Switzerland AG, AR9640) at 100°C for 10min. After additional washes, tissues were exposed to Novocastra™ Peroxidase Block solution (Leica Biosystems, RE7101) for 10min. Sections were incubated with appropriate primary antibody dilutions for 60 minutes at room temperature followed by washes with BOND Wash Solution 1X (BOND Wash Solution 10X Concentrate, Biosystems Switzerland AG, AR9590). Next, tissues were incubated for 60 minutes with HRP-conjugated secondary antibody. After washes in BOND Wash Solution, the sections were first exposed to Opal 570 fluorophore for 10min. BOND Epitope Retrieval Solution 1, pH6 (Biosystems Switzerland AG, AR9961) was applied to the tissues for 15min at 95°C to prepare the second cycle of hybridization, starting again with washes, blocking, primary and secondary antibody. Next the sections were exposed to Opal 690 fluorophore for 10min. After a final incubation in BOND Epitope Retrieval Solution 1, pH6, the sections were washed and incubated with Spectral DAPI 1X (10X Spectral DAPI, FP1490, Akoya Biosciences). Last, sections were washed in DI water. Anti-Fade Fluorescence Mounting Medium (abcam, ab104135) and Coverslips 24×50mm #1,5 (Epredia, BB02400500SC13MNZ0) were used to mount the tissue prior to imaging with Stellaris 5 microscope (Leica). Antibodies dilutions are listed in Table S5.

#### Transmission Electron Microscopy

Apical-in organoids were removed from Matrigel using Cell Recovery solution (Corning, 354253) for 50 min maximum on ice and then washed with cold PBS pH (7.4) (GIBCO, 10010-015). Apical-out organoids (not damaged and damaged) were collected by centrifugation. TOs were then fixed were immersed in a fixation solution containing 2% paraformaldehyde (Electron Microscopy Science, 15700) and 2.5% glutaraldehyde (Electron Microscopy Science, 16300) in 0.1M Cacodylate pH 7.4 buffer (Sigma-Aldrich, C0250) for 1 hour at room temperature and then overnight at 4°C. After five washes with 0.1 M cacodylate buffer (pH 7.4), the samples were post-fixed for 1 hour in a solution containing 1.5% potassium ferrocyanide and 1% osmium tetroxide in 0.1 M cacodylate buffer (pH7.4). The samples were washed with bi-distillated water and then incubated with 1% uranyl acetate for 20 minutes. After five washes in ddH2O and dehydration steps in graded alcohol series, the samples were embedded in epon resin for 12 h and polymerized at 60°C for 48hrs. For transmission electron microscopy (TEM) analysis, a region of interest was selected under light. After trimming, silver/gray thin sections (50 nm thickness) were collected on formvar-coated single-slot copper grids. After post-staining with 1% uranyl acetate and lead citrate (Electron Microscopy Science, 22410-01), 5 min each, images were recorded using a FEI Tecnai Spirit (FEI Company) operated at 120 keV using a side-mounted 2K × 2K CCD camera (Veleta, Olympus).

#### Scanning Electron Microscopy

Apical-in organoids were removed from Matrigel using Cell Recovery solution (Corning, 354253) for 50 min maximum on ice and then washed with cold PBS pH (7.4) (GIBCO, 10010-015). Apical-out organoids (not damaged and damaged) were collected by centrifugation. TOs were then immersed in a fixation solution containing 2.5% glutaraldehyde (Electron Microscopy Science, 16200) and 2% paraformaldehyde (Electron Microscopy Science, 15710) in 0.1 M cacodylate buffer pH 7.4 for 1 hour and then overnight at 4°C. After three washes with 0.1M cacodylate buffer pH7.4, the samples were incubated in 1% tannic acid (Sigma-Aldrich, 21700) in 0.1M cacodylate buffer (pH7.4). After another round of three washes with 0.1 M cacodylate buffer (pH7.4) the samples were incubated in 1% osmium in 0.1 M cacodylate buffer (pH7.4). The samples were rinsed with bi-distillated water, dehydrated in graded ethanol series and dried with hexamethyldisilazane (Sigma-Aldrich, 379212). Additionally, the organoid preparations were dried in an oven at 60°C for 5 min. and mounted on aluminum stubs by depositing them on double-sided carbon tape and sputter coated (Quorum; SC7620) with gold/palladium (5–8 nm). The organoids were imaged at 4kV and 200pA with a scanning electron microscope (SEM Merlin; Zeiss) using the HE-SE2 detector.

#### Mass spectrometry of conditioned media

Conditioned media collected after 24h from trophoblast organoids culture was stored at −80°C and then thawed on ice. Protein concentrations of the media and secretome samples were measured to be between 2-3 g/l using a NanoDrop spectrophotometer (Thermo Scientific). Aliquots of 200 μL were taken from each sample (app. 500 μg per sample), and proteins were denatured in 1% sodium deoxycholate (SDC) in 50 mM HEPES buffer, pH 8.5. To ensure complete protein solubilization, samples were sonicated using a Branson digital sonifier with a tapered microtip using 70% output for 10 s on ice. After reduction and alkylation in 2 mM tris-(2-Carboxyethyl)phosphine hydrochloride (TCEP) and 10 mM chloroacetamide at room temperature for 60 min in the dark, proteins were digested with TPCK treated trypsin (Thermo Fisher Scientific) and Lys-C Fujifilm Wako at an enzyme to protein ratio of 1:100 and 1:500, respectively, over night at 37°C, with shaking. On the next day, an additional aliquot of trypsin (1:100) was added for 3 h. For TMT peptide labeling, 9 channels of a TMT 10plex reagent kit (Thermo Fisher Scientific, lot# TI273434), 126 – 130C, were used according to the manufacturer’s instructions. In short, TMT reagents were dissolved in 100 μL of anhydrous acetonitrile, then mixed with their respective sample in 70 μL, pH adjusted to pH 8.5 using 50 μL 1M HEPES, pH 8.5, and incubated for 1h at room temperature in the dark with shaking. The reaction was quenched with the addition of 2 μL 5% Hydroxylamine solution, the samples were pooled, speed-vacuumed for 30 min and desalted using a Sep-Pak® Vac tC18 cartridge 1 cc/50 mg (Waters). In brief, the Sep-Pak cartridge was conditioned with 100% methanol and equilibrated using 0.15% TFA in water. After loading the TMT peptide sample pool, the Sep-Pak cartridge was washed 3x with 0.15% TFA in water, peptides were eluted with 50% acetonitrile solution in water and 0.15% TFA, then dried in a speed vac. TMT peptides were separated using a gradient composed of 20 mM ammonium formate in either water (buffer A) or in 90% acetonitrile with water (buffer B) at pH 10 on a reversed-phase YMC Triart C18 0.5 × 250 mm column (YMC Europe GmbH), concatenated into 48 “super-fractions”, dried, and each fraction was reconstituted with 15 μl of 2% acetonitrile in water with 0.1% TFA. Approximately 1 μg per sample was analyzed by LC-MS/MS using a Vanquish Neo chromatography system with a trap-and-elute configuration, connected to an EASY-Spray source and an Orbitrap Eclipse Tribrid MS (all Thermo Fisher Scientific). The built-in Comet search engine identified TMT-labeled peptides in real-time from MS2, prompting MS3 scans of selected precursors for subsequent quantification.

#### RNA extraction

Apical-in TOs were collected and Matrigel was removed using Cell Recovery Solution (Corning, 354253) for 50 min maximum on ice followed by one wash with cold PBS pH (7.4) (GIBCO, 10010-015). Apical-out TOs were directly collected from the well followed by one wash with cold PBS pH (7.4) (GIBCO, 10010-015). Total RNA was extracted from organoid pellets using the RNeasy Micro kit with on column DNase treatment (Qiagen, 76004), following manufacturer’s instructions. The RNA was resuspended in 14 μl RNase-free water. The purity and concentration of the RNA was determined using the UV-Vis Spectrophotometer NP80 (Implen).

#### cDNA synthesis

For cDNA synthesis, 300 ng-1 μg of total RNA was reverse transcribed using SuperScript VILO cDNA Synthesis Kit (Thermo Fisher Scientific, 11754050) following manufacturer’s instructions. The extracted RNA was diluted into 5X VILO Reaction Mix containing random primers, dNTPs, and MgCl2 and 10X SuperScript III Enzyme Blend containing SuperScript III Reverse Transcriptase, RNase Recombinant Ribonuclease Inhibitor, and proprietary helper protein. The samples were then incubated for 10 minutes at 25°C, 1 hour at 42°C and 5 minutes at 85°C. An RNA sample without Reverse transcriptase was used as control for genomic DNA contamination.

#### Real-time quantitative PCR (RT-qPCR)

RT-qPCR was performed with the StepOnePlus PCR system (Applied Biosystems) using TaqMan Fast Advanced Master Mix (Thermo Fisher Scientific, 4444557) and Taqman gene specific primer probes (*ERVW-1*, Hs01926764; *GCM1*, HS00961602; *MKI67*, HS00606991; *CGB3*, Hs00361224; *PCNA*, Hs00427214; *GJA1*, Hs00748445; *SDC1*, Hs00896423), following manufacturer’s protocol. The cycling conditions followed were 20 seconds at 95°C and 40 cycles of 3 seconds at 95°C followed by 30 seconds at 60°C.

#### hCG ELISA

Supernatants and organoids pellet were collected and stored at −80°C. Dilution factors were first determined on test samples before final experimentation. Manufacturer’s instructions were followed in the execution of the ELISAs.

### QUANTIFICATION AND STATISTICAL ANALYSIS

#### SYN volume quantification

CDH1 and SDC1 signals were analysed and combined to get the TOs total volume using ImageJ software, whereas SYN volume has been determined subtracting the total volume from the CDH1 (VCT) volume. SYN volume was then plotted as percentage of the total organoid volume. Statistical significance was determined using t-test. Significance was determined by p<0.05. ****p<0.0001.

#### qPCR data analysis

Expression levels were calculated using the comparative Cycle threshold (Ct) method. The geometric mean of *HPRT1*, *TOP1*, and *TBP* housekeeping genes was used for normalization of the relative gene expression levels. Normalized expression levels were calculated as 2^-ΔCt^ where ΔCt= Ct_gene of interest_ – Ct_geometric mean of housekeeping genes_. Each qPCR reaction was performed in duplicate, and a non-template control was always included. Statistical significance has been determined as stated in figure legend. Ns = not significant, or p-value above 0.05. *p<0.05, **p<0.01.

#### hCG ELISA data analyses

Graphpad Prism was used to create visual graphs and analyses. Statistical significance was determined by t-test. Ns = not significant. Normalization was performed using RNA concentration.

#### Bulk RNA-seq analysis

Paired-end sequence reads were aligned to the human genome (GRCh38 primary assembly) using the"qAlign" function from the Bioconductor package QuasR version 1.48.1 [53] with default parameters except for aligner = "Rhisat2", splicedAlignment = "TRUE", allowing only uniquely mapping reads. Raw gene counts were generated using the "qCount" function (QuasR) with GENCODE release 38 ‘basic gene annotation’ ("gencode.v38.annotation.gtf" as TxDb object) as query, with default parameters except useRead=first and orientation=opposite.The count table was modified by replacing the Ensembl gene ID with the gene symbol (if unique) or a combination of the gene symbol and the Ensembl ID (if the gene symbol was not unique) or keeping the Ensembl ID (for genes without a gene symbol) as the unique identifier. Genes were classified as significantly upregulated or downregulated based on the threshold stated in figure legends. GO analysis have been performed using Enrichr [51]. All source code used are available from GitHub: https://github.com/fmicompbio/apical-out-placental-organoids.

#### *TNFRSF12A* expression in PAMM

To examine the expression of *TNFRSF12A* in cell populations of PAMM (also known as PAMM1a) and intervillous monocytes (ivMonos, also known as PAMM1b), bulk RNA-seq data profiling of these cells [54] were downloaded from EMBL-EBI ArrayExpress (accession number E-MTAB-14088). Normalized expression values (RPKM) were used to compare TWEAK expressions between the two populations.

#### Secretome analysis

MS2 spectra were searched using Sequest within Proteome Discoverer against the HUMAN Uniprot database and common contaminants. PSMs were validated to 1% FDR using a target-decoy strategy with Percolator. Quantification was based on MS3 TMT reporter ions, requiring minimum SPS match of 65%, with no correction applied to quant values. Protein-level data were processed using the einprot R package (v0.9.7) [55]. Contaminants were removed, and only high-confidence Master proteins (score >2, ≥2 peptides) were retained. Intensities were log2-transformed, normalized (center.median), and imputated (MinProb). Differential abundance was assessed using limma [50, 56], and enrichment on pathway-level, in Gene Ontology terms, and in known complexes was tested using camera [57]. Significance thresholds were set at adjusted p-value <0.02 and absolute log2 fold changes ≥ 2.

## Acknowledgments

We thank G. Burton and A. Moffett for insightful feedback on the manuscript. We are grateful to the Turco laboratory for feedback and fruitful discussions. Thank you to the members of the Facility for Advanced Imaging and Microscopy, L. Gelman, L. Plantard, J. Eglinger, and T.-O. Buchholz for their support with imaging and quantification. We thank the functional genomics facility for performing RNA sequencing experiments. We are grateful to Holly Anderson for the derivation of trophoblast organoids. We thank Emy Bosseboeuf and Caitlin Duncan (University of Cambridge) for isolating the primary monocytes and to patients for their donations. This work was supported by the by the Swiss National Science Foundation (SNSF) (grant number 10004677) and the Novartis Research Foundation. Schemes were created in https://BioRender.com

## Authors contributions

I.C. and M.Y.T. conceived and designed experiments. I.C., G.G., L.K. performed experiments. A.G.M performed TEM and SEM images. J.S. performed LC-MS/MS. I.C., G.G., J.S., H.R.H. analyzed data. I.C. and M.Y.T. interpreted the data. I.C. and M.Y.T. wrote the manuscript. Q. L. performed TWEAK gene expression analysis. N.M. provided primary monocytes. I.C, G.G., A.G.M., L.K., J.S., Q. L., H.R.H., N.M., M.Y.T. reviewed the manuscript.

---

**Figure S1.**
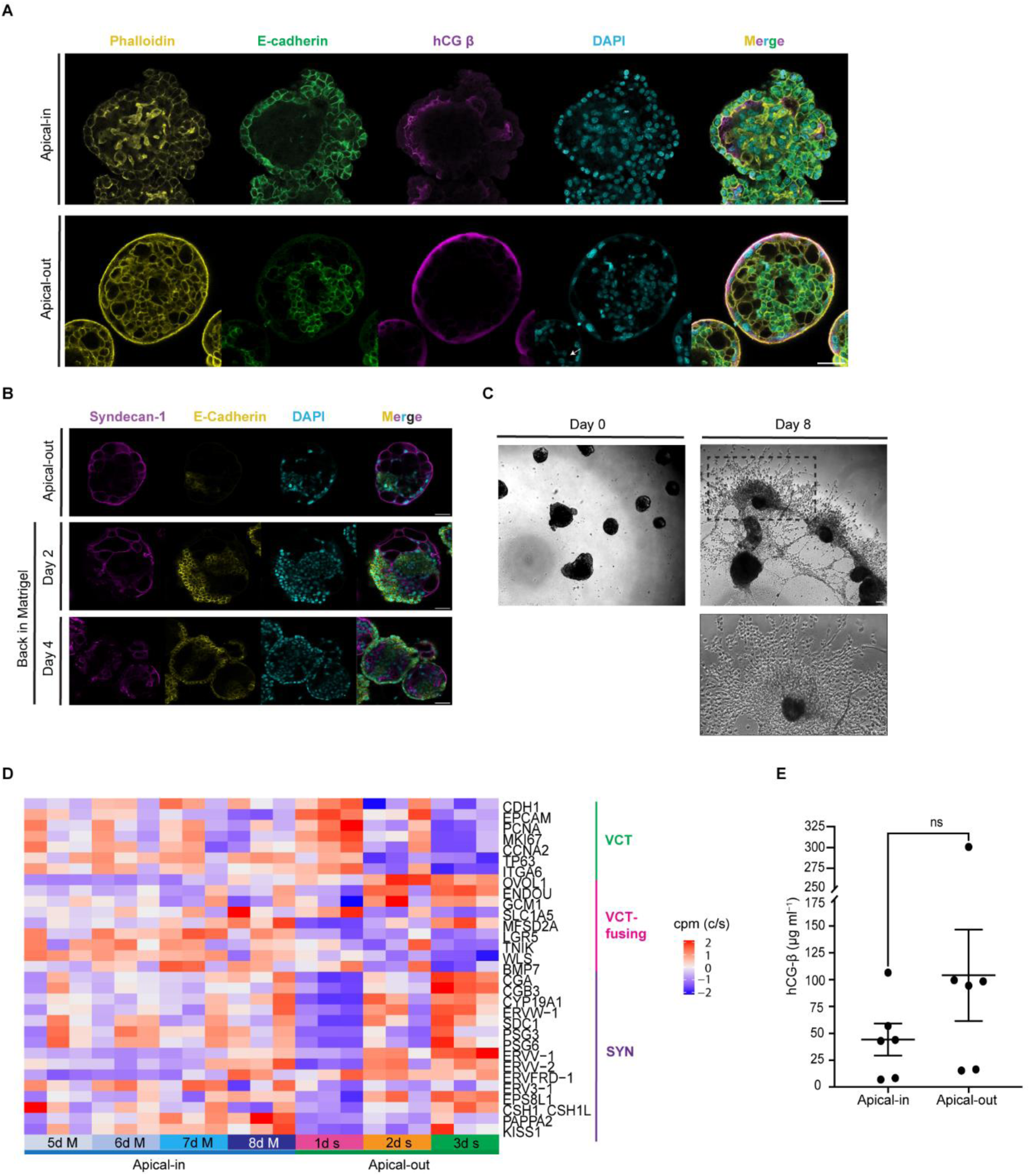
A: Confocal microscopy images of apical-in and apical-out trophoblast organoid stained for Phalloidin, hCG β, E-cadherin and DAPI (representative image from *n* = 8 and *n* = 7 respectively). Scale bars, 50 μm. B: Confocal microscopy images of apical-out (day 3 in suspension culture) and apical-out back in Matrigel (day 2 and day 4) trophoblast organoids stained for E-cadherin, syndecan-1 and DAPI with merged images (representative image from *n*=5, *n*=7 and *n*=4 respectively). Scale bars, 50 μm. C: Representative bright-field images of apical-out trophoblast organoid in EVT medium at day 0 and at day 8. Scale bars,100 μm. Data from three independent organoid lines (X113, Y060, and Z143). D: Heatmap depicting centered and scaled counts per million (cpm) of selected genes expressed in apical-in and apical-out trophoblast organoids over a time course experiment and associated with markers of villous cytotrophoblast (VCT), VCT-fusing and syncytiotrophoblast (SYN). E: hCG-β ELISA from conditioned medium of apical-in and apical-out trophoblast organoids. Mean values ± SEM. hCG-β amount (μg ml^−1^) secreted in 24h (between days 7 and 8 in Matrigel and days 2 and 3 in suspension). Data are normalized to RNA concentration. The P value was determined using paired t test. ns, not significant. N = 6 independent experiments. Data from three independent donors (X113, Y031, and X087).

**Figure S2.**
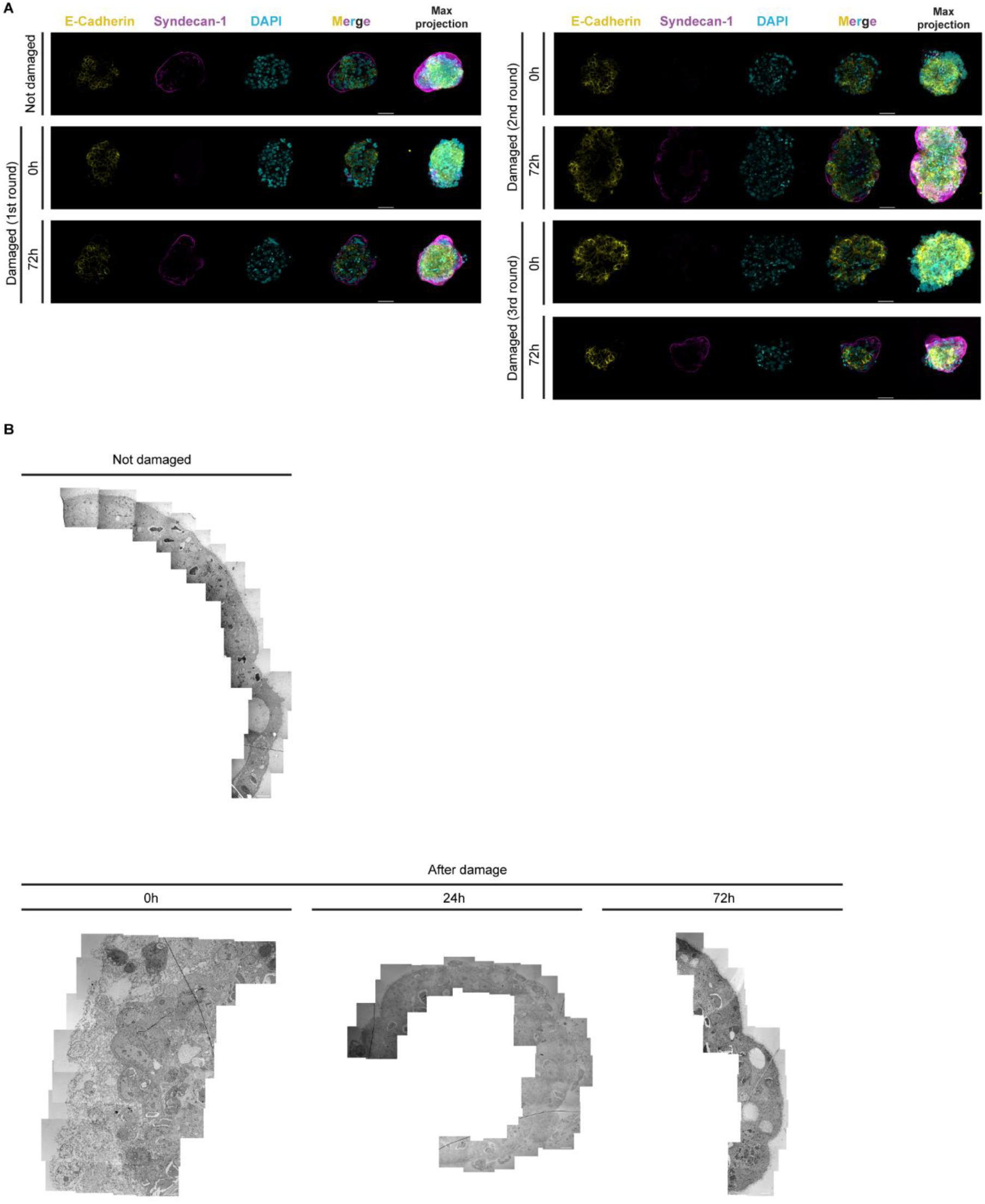
A: Confocal microscopy images of before damage and damaged TOs (subjected to the damage protocol three times) stained for E-cadherin, syndecan-1 and DAPI, with merged images and max projection. Scale bars, 50 μm. B: Representative images after performing stitching of Transmission Electron Microscopy time-course images of apical-out trophoblast organoids before damage and after damage (0h, 24h and 72h). Scale bars are 5 µm.

**Figure S3.**
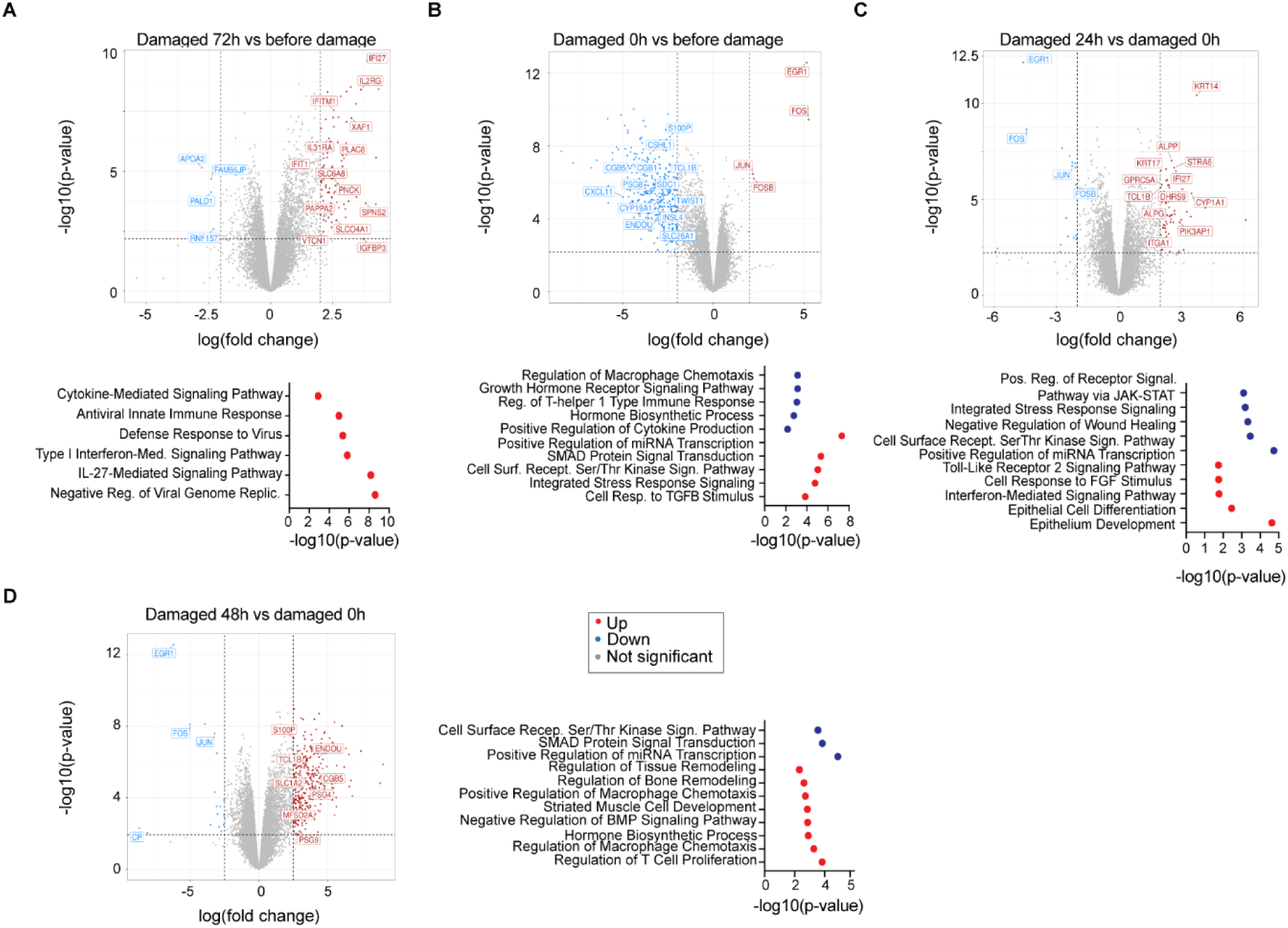
A: On the top panel, volcano plot highlighting selected DEGs from the comparison between damaged organoids at time point 72h vs before damage (upregulated genes: 94, downregulated genes: 6, not significant: 15604; abs(logFC) > 2, logCPM > 1, FDR < 0.05). Below, biological processes upregulated in damaged organoids at time point 72h. B: On the top, volcano plot highlighting selected DEGs from the comparison between damaged organoids at time point 0h vs before damage organoids (upregulated genes: 4, downregulated genes: 404, not significant: 15296; abs(logFC) > 2, logCPM > 1, FDR < 0.05). Below, biological processes downregulated (in blue) and upregulated (in red) in damaged organoids at time point 0h. C: On the top panels, volcano plot highlighting selected DEGs from the comparison between damaged organoids at time point 24h vs 0h (upregulated genes: 72, downregulated genes: 22, not significant: 15610). Below, biological processes downregulated (in blue) and upregulated (in red) in damaged organoids at time point 24h. D: On the left, volcano plot highlighting selected DEGs from the comparison between damaged organoids at time point 48h vs 0h (upregulated genes: 369, downregulated genes: 20, not significant 15315; abs(logFC) > 2, logCPM > 1, FDR < 0.05). on the right, biological processes downregulated (in blue) and upregulated (in red) in damaged organoids at time point 48 h. Data from three independent donors (X113, Y031, and Y060).

**Figure S4.**
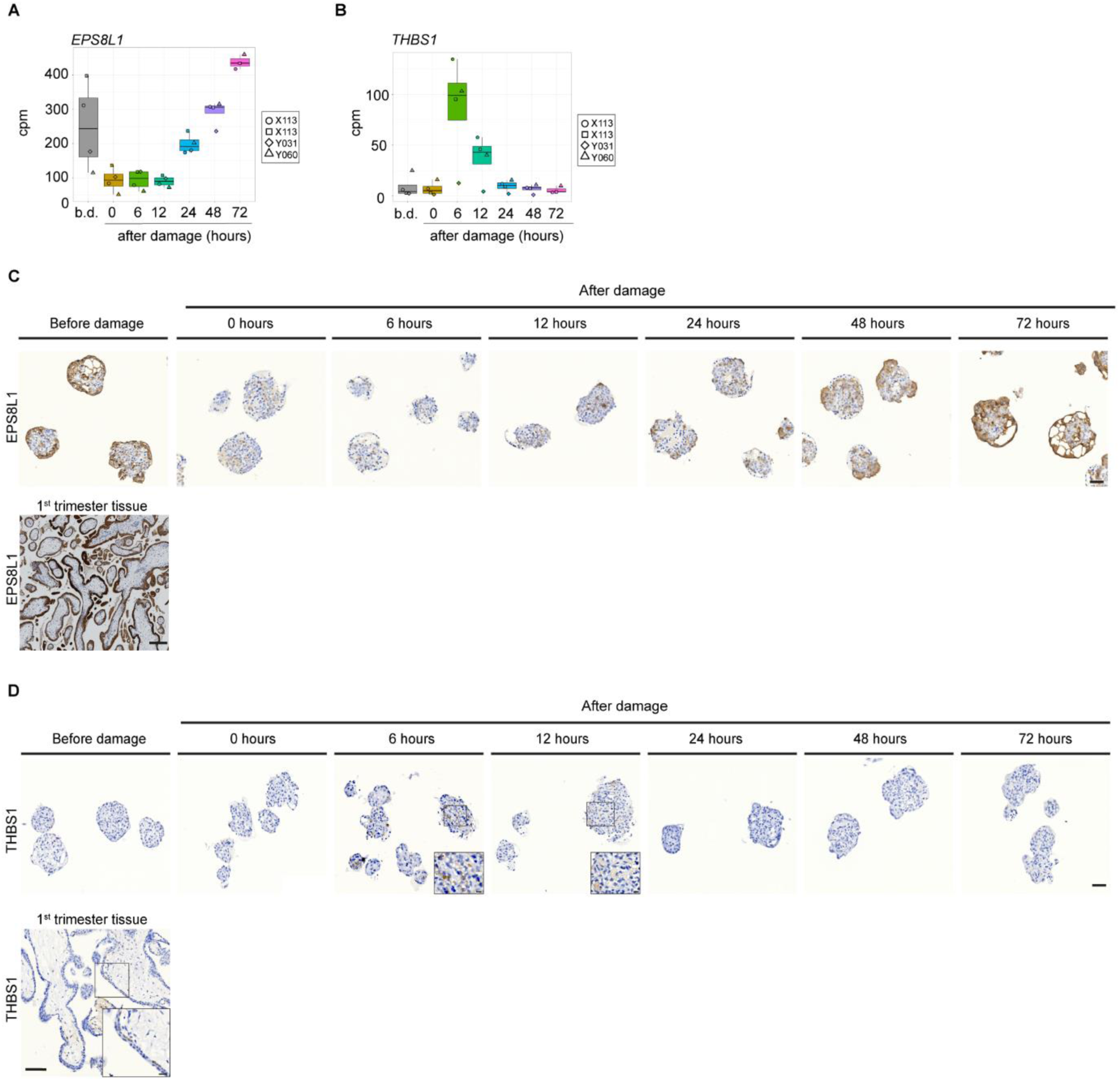
A: Expression level (in cpm) of *EPS8L1* across the time-course experiment. B: Expression level (in cpm) of *THBS1* across the time-course experiment. C: Representative images of EPS8L1 staining across the time-course experiment in TOs (scale bars, 50 μm) and in first trimester placenta tissue section (scale bars, 200 μm). Data from three independent donors. D: Representative images of THBS1 staining across the time-course experiment in TOs (scale bars, 50 μm, magnified box scale bars, 10 μm) and in first trimester placenta tissue section (scale bars, 100 μm; magnified box scale bars, 20 μm). Data from three independent donors.

**Figure S5.**
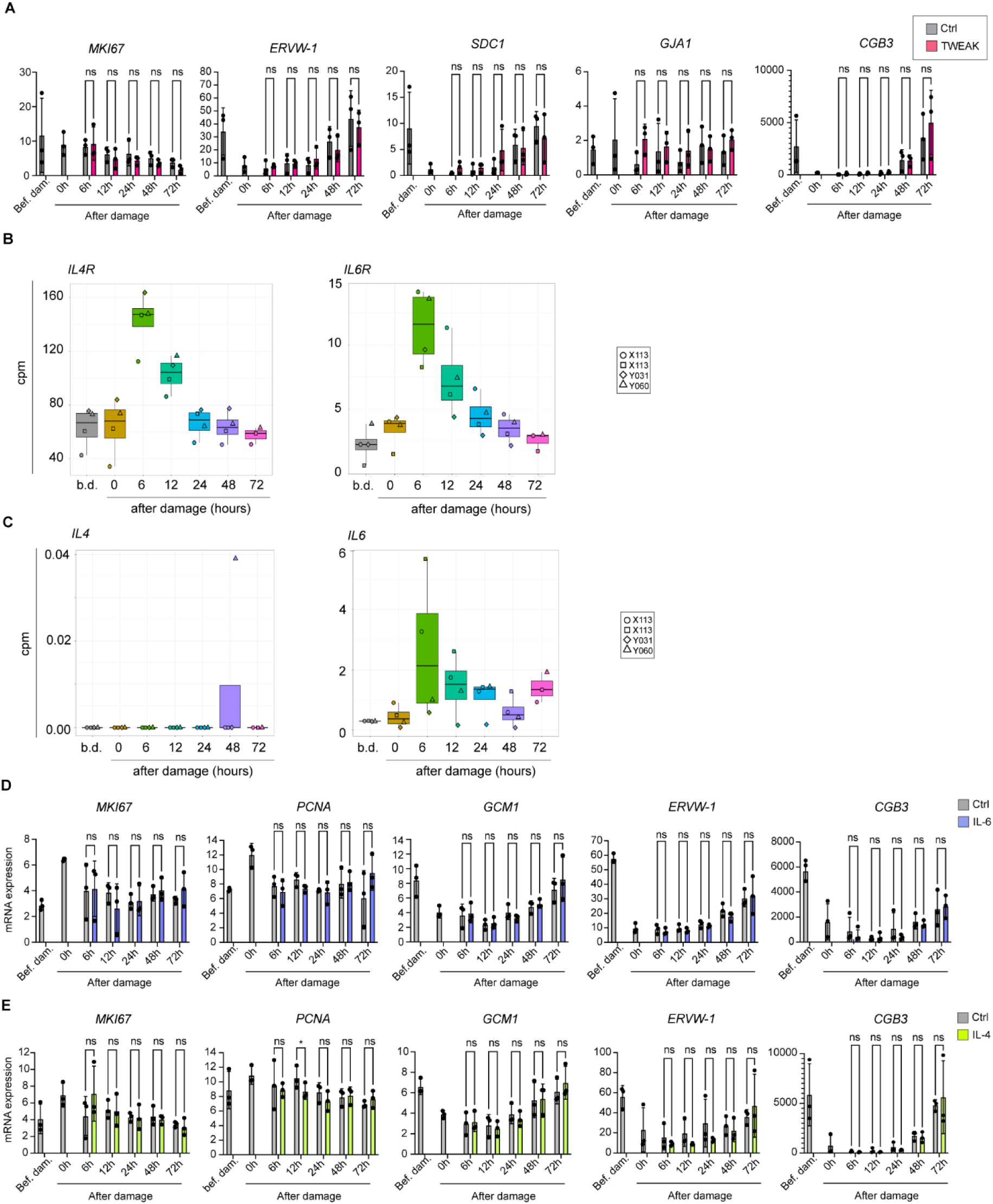
A: qPCR showing mRNA expression of *MKI67, ERVW-1, SDC1, GJA1,* and *CGB3* in control and TWEAK-treated TOs. Mean values ± SD measured in duplicates (normalized to housekeeping genes, *TBP, TOP1* and *HPRT1*) are shown. The P values were determined using multiple comparison test. ns, not significant; Data from three independent experiments. B: Expression level (in cpm) of *IL4R* and *IL6R* gene across the time-course experiment. C: Expression level (in cpm) of *IL4* and *IL6* gene across the time-course experiment. D: qPCR showing mRNA expression of *MKI67, PCNA, GCM1, ERVW-1, and CGB3* in control and IL-6-treated TOs. Mean values ± SD measured in duplicates (normalized to housekeeping genes, *TBP, TOP1* and *HPRT1*) are shown. The P values were determined using multiple comparison test. ns, not significant. Data from three independent experiments. E: qPCR showing mRNA expression of *MKI67, PCNA, GCM1, ERVW-1, and CGB3* in control and IL-4-treated TOs. Mean values ± SD measured in duplicates (normalized to housekeeping genes, *TBP, TOP1* and *HPRT1*) are shown. The P values were determined using multiple comparison test. ns, not significant; * p < 0.05. Data from three independent experiments.

**Table S1**

List of genes from bulk RNA sequencing comparing apical-in and apical-out trophoblast organoids.

**Table S2**

LC-MS/MS analysis of trophoblast organoids supernatants.

**Table S3**

List of genes from bulk RNA sequencing comparing trophoblast organoids over the damage time-course experiment.

**Table S4**

Trophoblast organoids donor information.

**Table S5**

List of antibodies used in immunofluorescence (IF) and immunohistochemistry (IHC).

**Movie S1**

3D reconstruction of apical-in trophoblast organoid stained for E-cadherin, syndecan-1 and DAPI. Scale bars, 50 μm.

**Movie S2**

3D reconstruction of apical-out trophoblast organoid stained for E-cadherin, syndecan-1 and DAPI. Scale bars, 50 μm.

